# Synaptotagmin 9 modulates spontaneous neurotransmitter release in striatal neurons by regulating substance P secretion

**DOI:** 10.1101/2022.04.18.488681

**Authors:** Michael J. Seibert, Chantell S. Evans, Kevin S. Stanley, Zhenyong Wu, Edwin R. Chapman

## Abstract

Synaptotagmin 9 (SYT9) is a tandem C2-domain Ca^2+^ sensor for exocytosis in neuroendocrine cells; its function in neurons remains unclear. Here, we show that endogenous SYT9 does not trigger rapid synaptic vesicle exocytosis in cultured cortical, hippocampal, or striatal neurons; rather, synaptotagmin 1 (SYT1) fulfills this function. SYT9 is able to regulate evoked synaptic vesicle exocytosis, but only when massively over-expressed. In striatal neurons, loss of SYT9 reduced the rate of spontaneous miniature neurotransmitter release events (minis). To delve into the underlying mechanism, we localized SYT9 within these neurons and found that it is targeted to dense core vesicles, where it regulates the release of substance P (SP), a neuropeptide known to modulate mini frequency. Exogenous SP, but not other striatal peptide hormones, completely rescued the *Syt9* KO mini phenotype. Biochemical experiments revealed that Ca^2+^-binding to the C2A domain of SYT9 triggers membrane fusion *in vitro*, and mutations that disrupt this activity abolished the ability of SYT9 to regulate both SP release and mini frequency in striatal neurons. We conclude that SYT9 indirectly regulates synaptic transmission in striatal neurons by controlling SP release.

## INTRODUCTION

The synaptotagmins (SYTs) are a large family of membrane trafficking proteins; seventeen isoforms have been identified in mammals, each encoded by a distinct gene (Craxton, 2010; Wolfes & Dean, 2020). In addition, most isoforms are thought to undergo alternative splicing, generating further potential diversity within this family (Craxton, 2010). Most isoforms have a single N-terminal transmembrane domain (TMD) with a short intra-luminal domain. In all isoforms, the cytoplasmic domain largely comprises tandem C2-domains that are connected by a short linker. In at least eight of the seventeen isoforms, the C2-domains serve as Ca^2+^-dependent anionic lipid-binding motifs (Bhalla et al., 2008; Rickman et al., 2004). Ca^2+^ binding is thought to be mediated by acidic side chains in two flexible loops that protrude from each C2-domain (Sutton et al., 1995). In the case of SYT1, Ca^2+^ triggers the partial penetration of these loops into lipid bilayers and facilitates interactions with SNARE proteins (Chapman, 2008). While membrane penetration is an essential step in Ca^2+^-triggered exocytosis (Bai et al., 2016; Evans et al., 2015; Liu et al., 2014), the role of SYT-SNARE interactions in membrane fusion remains unresolved (Bai et al., 2016; Bhalla et al., 2006; Chicka et al., 2008; Das et al., 2020; Van Den Bogaart et al., 2011; Zhou et al., 2017).

A current goal is to understand the functional diversity within the SYT family of proteins. SYT1 and SYT2 have been heavily studied and shown to serve as Ca^2+^ sensors that trigger rapid, synchronous synaptic vesicle (SV) exocytosis in neurons (Geppert et al., 1994; Pang, Melicoff, et al., 2006; Stevens & Sullivan, 2003). SYT1 also regulates SV docking (Chang et al., 2018; Liu et al., 2009; Reist et al., 1998), endocytosis (Jorgensen et al., 1995; Nicholson-Tomishima & Ryan, 2004; Poskanzer et al., 2003; Yao et al., 2012), and spontaneous release (Bai et al., 2016; Courtney et al., 2019; DiAntonio & Schwarz, 1994; Kerr et al., 2008; Littleton et al., 1994; Vevea & Chapman, 2020). Importantly, SYT1 and SYT2 form a closely related clade with one more family member, SYT9 (Rickman et al., 2004). This homology prompted the idea that SYT9 might also function as a fast Ca^2+^ sensor for SV release, which is a point we address in detail in the current study. For clarity, we note that SYT9 is sometimes referred to as SYT5, and vice versa; the SYT9 studied here comprises 386 residues (NCBI Gene accession # 53420) whereas SYT5 is 491 residues in length (NCBI Gene accession #60510).

Along with SYT1, SYT9 co-regulates catecholamine secretion from dense core vesicles (DCV) in PC12 cells (Fukuda et al., 2002; Lynch & Martin, 2007). SYT9 also regulates the release of follicle-stimulating hormone (FSH) from DCVs in the gonadotrophs of female mice in a sex-specific manner (Roper et al., 2015). However, while SYT9 is widely expressed in the brain ([Mittelsteadt et al., 2009]; note: these authors referred to SYT9 as SYT5), its function in neurons remains controversial. One report concluded that SYT9 serves as a Ca^2+^ sensor for rapid SV exocytosis in cultured striatal neurons (Xu et al., 2007). In contrast, SYT9 failed to support SV exocytosis in *Syt1* KO hippocampal neurons, and pHluorin-tagged SYT9 underwent regulated exocytosis in both axons and dendrites of hippocampal neurons, with kinetics suggestive of DCV fusion (Dean, Dunning, et al., 2012).

To address and clarify the function of SYT9 in neurons, we studied synaptic transmission in cultured hippocampal, cortical, and striatal neurons obtained from *Syt9* KO mice; for completeness *Syt1* KO neurons were examined in parallel. We show that all three classes of neurons express both SYT9 and SYT1, but loss of SYT9 had no discernable effect on evoked neurotransmitter release, while loss of SYT1 disrupted rapid transmission in all three neuronal preparations. SYT9 was able to rescue the loss of synchronous release that is characteristic of *Syt1* KO neurons, but only when over-expressed by ∼25-fold, which matched the expression levels of endogenous SYT1. Interestingly, loss of SYT9 in cultured striatal neurons resulted in a specific and unexpected phenotype: a decrease in the spontaneous miniature SV fusion rate (i.e., mini frequency). We then discovered that SYT9 was co-localized with substance P (SP), a DCV peptide hormone known to be enriched in striatal neurons. Moreover, loss of SYT9 impaired action potential-triggered SP release. Importantly, in patch-clamp experiments, supplementing the bath with this neuropeptide rescued the mini phenotype. Moreover, the structural elements of Syt9 that underlie Ca^2+^ triggered membrane fusion *in vitro* also support evoked SP release and spontaneous SV fusion in living striatal neurons. These findings reveal a mechanism in which SYT9 indirectly regulates spontaneous transmission in striatal neurons by directly regulating secretion of the neuromodulator, SP.

## RESULTS

### Synaptotagmin 9 is not required for fast, synchronous neurotransmitter release in cultured neurons

Immunoblot analysis of neuronal culture lysates revealed that SYT1 and SYT9 are expressed in cortical, hippocampal, and striatal neurons (**Fig. 1a,e**). By including known amounts of purified recombinant SYT1 cytoplasmic domain (C2AB) or SYT9 (full-length) as standards on these blots we were able to determine the absolute expression levels via quantitative densitometry (**Fig. 1b,f**). This analysis revealed that SYT1 and SYT9 were ∼0.1% and ∼0.004% of the total protein in each of the neuronal lysates, respectively. Hence, SYT1 expression is ∼25 times greater than SYT9 in all three classes of neurons.

**Figure 1.**
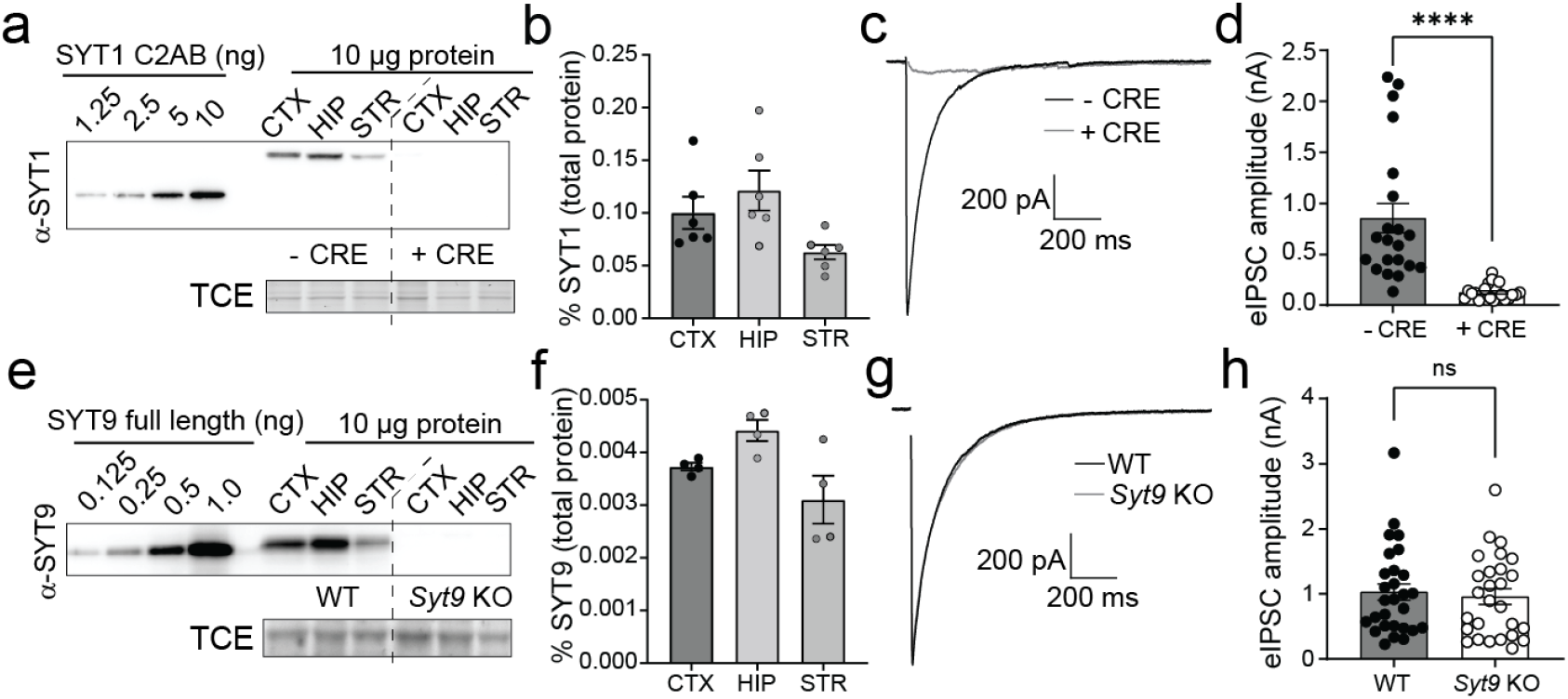
SYT9 is expressed in cultured cortical, hippocampal, and striatal neurons but does not support evoked neurotransmitter release. **a**, Immunoblot of day *in vitro* (DIV) 14 cortical (CTX), hippocampal (HIP), and striatal (STR) neuronal lysates from *Syt1* cKO neurons +/- CRE virus; recombinant SYT1 C2AB served as a standard to quantify SYT1 expression levels. TCE staining served as a loading control. **b**, Quantification of the SYT1 expression levels in each of the three neuronal lysates, as a fraction of total protein (n = 6). **c**, Averaged evoked IPSC (eIPSC) traces from *Syt1* cKO striatal neurons +/- CRE, and (**d**) peak eIPSC amplitudes from - CRE (n = 21) or *+* CRE (n = 22) cells, showing loss of synchronous release upon loss of SYT1 (p < 0.0001; Mann-Whitney test). **e**, Same as panel (a) but blotting for SYT9 in lysates from WT and *Syt9* KO striatal neurons, using the indicated amounts of a full-length recombinant SYT9 standard. **f**, Same as panel (**b**), but for SYT9 expression in WT and *Syt9* KO striatal neurons (n = 4); SYT9 is expressed at ∼25-fold lower levels than SYT1. **g-h**, Same as panels (**c-d**), but in WT (n = 28) and *Syt9* KO (n = 27) striatal neurons. In sharp contrast to *Syt1* cKO + CRE neurons, loss of SYT9 did not affect single eIPSCs in striatal neurons (p = 0.6979; unpaired Student’s t-test). ****p < 0.0001; ns indicates p > 0.05, and error bars represent SEM.

A prior study suggested that SYT1 and SYT9 serve as partially redundant Ca^2+^ sensors in striatal neurons and that SYT9 can fully rescue fast release in *Syt1* KO cortical neurons (Xu et al., 2007). However, this is difficult to reconcile with the well-established findings that loss of SYT1 in hippocampal and cortical neurons disrupts virtually all synchronous release (Geppert et al., 1994; Maximov & Südhof, 2005). If SYT9 is a sensor for synchronous release, some fast release should have persisted in the *Syt1* KO neurons. To address this conundrum, whole-cell patch-clamp experiments were conducted using cultured neurons from *Syt1* conditional KO (cKO) and *Syt9* KO mice. In striatal *Syt1* cKO neurons, ablation of SYT1 expression with application of CRE lentivirus eliminated fast, synchronous neurotransmitter release with an 8-fold reduction in evoked inhibitory postsynaptic current (eIPSC) amplitude (**Fig. 1c,d**). Disruption of SYT9 expression, however, did not result in any change in eIPSC amplitude or kinetics in striatal neurons (**Fig. 1g,h**). Analogous results were obtained using cultured cortical and hippocampal neurons (**Supp. Fig. 1**). Evoked release during a stimulus train was also unaffected in *Syt9* KO cortical, hippocampal, and striatal neurons, further demonstrating that this isoform does not impact evoked transmission (**Supp. Fig. 2**). We conclude that endogenous SYT9 is unable to support synchronous release in absence of SYT1 in any of the neuronal preparations studied here, perhaps because of the relatively low expression levels of this protein.

### SYT9 rescues SYT1 KO phenotypes only when over-expressed

Xu et al. (2007) reported that SYT9 over-expression can functionally rescue synchronous neurotransmitter release in cultured *Syt1* KO cortical neurons (Xu et al., 2007). Similar observations were made upon over-expression of SYT9 in a calyx of Held slice preparation lacking the primary Ca^2+^ sensor at this synapse, SYT2 (Kochubey et al., 2016). Given the observation that endogenous SYT9 is expressed at levels ∼25-fold lower than SYT1 (**Fig. 1b,f**), a quantitative approach was taken to re-evaluate this previously reported rescue activity. Cultured *Syt1* cKO cortical neurons were treated at DIV1 with a CRE lentivirus to disrupt SYT1 expression and then transduced at DIV5 with lentivirus to express SYT9 with an N-terminal pHluorin tag (pH-SYT9). The pH-SYT9 lentivirus was titered such that expression of pH-SYT9 in the + CRE neurons would be either 1x or 25x the endogenous SYT9 expression level; the latter condition matching the levels of endogenous SYT1 (**Fig. 2a**; **Fig. 1**). Ablation of SYT1 expression again resulted in a loss of fast, synchronous neurotransmission, and 1x over-expression of pH-SYT9 had no significant effect on eIPSC amplitude, and while there was a decrease in the time-to-peak this did not reach significance (**Fig. 2b-d**). However, 25x over-expression of pH-SYT9 significantly rescued the eIPSC amplitude (60% compared to - CRE) and completely rescued the eIPSC time-to-peak (**Fig. 2b-d**). The 25x over-expression condition also rescued the decay time of the + CRE condition (**Supp. Fig. 3**). So, only supraphysiological SYT9 expression levels rescue evoked responses in neurons lacking SYT1 (Kochubey et al., 2016; Xu et al., 2007).

**Figure 2.**
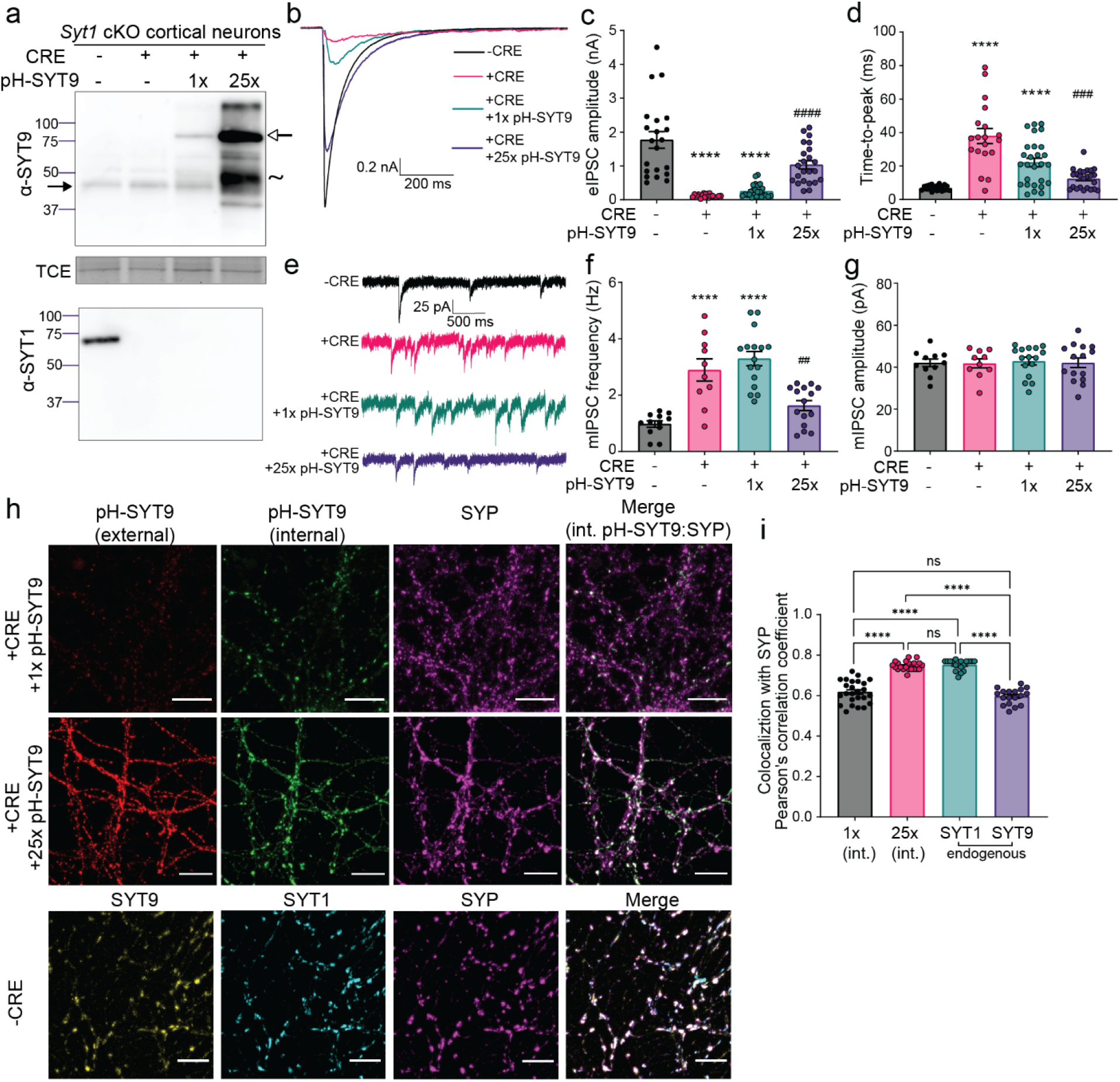
SYT9 rescues SYT1 KO phenotypes only when over-expressed. **a**, Immunoblots of DIV14 cortical neuronal lysates from *Syt1* cKO mice, with or without addition of CRE lentivirus. TCE staining served as a loading control (note: in all trials the α-SYT9 blot was stripped and re-probed with an α-SYT1 antibody, so TCE control is shown only once). + CRE neurons were later transduced with lentivirus encoding pHluorin-tagged SYT9 (pH-SYT9), resulting in either a 1x or 25x degree of over-expression as compared to endogenous SYT9. The closed arrowhead indicates endogenous SYT9, the open arrowhead indicates pH-SYT9, and the ∼ indicates proteolytic fragments from pH-SYT9. Averaged eIPSC traces (**b**), amplitudes (**c**), and time-to-peak for each condition (**d**), are plotted for - CRE (n = 21), + CRE (n = 19), + CRE +1x pH-SYT9 (n = 25), and + CRE +25x pH-SYT9 (n = 23) conditions. SYT9 partially rescued eIPSC amplitude and time-to-peak, but only when over-expressed 25-fold. **e**, Representative miniature IPSC (mIPSC) traces, showing increased rates of mIPSCs in *Syt1* cKO + CRE (n = 10) neurons compared to - CRE (n = 12). 1x over-expression of SYT9 was without effect (n = 16); full rescue of the unclamped phenotype was observed at 25x over-expression (n = 15); these data are quantified in (**f**). **g**, mIPSC amplitude was unaffected under all conditions. **h**, Representative immunofluorescence images of cultured *Syt1* cKO cortical neurons +/- CRE, with and without over-expressed pH-SYT9. Rescue constructs were tagged at the N-terminus with a pHluorin to enable differential labeling of the surface and internal fractions using α-GFP antibodies, see Methods for details. pH-SYT9 was present in both the plasma membrane (red) and on internal compartments (green). SVs were marked using an α-synaptophysin (SYP) antibody (magenta). **i**, Quantification of the colocalization between synaptophysin and: internal pH-SYT9, endogenous SYT1, and endogenous SYT9. In **c**-**g**, asterisks (*) indicate statistical comparisons (one-way ANOVA with Šídák’s multiple comparisons test) to *Syt1* cKO neurons without CRE, where ****p < 0.0001; hash symbols (#) indicate statistical comparisons to *Syt1* cKO neurons + CRE, where ##p < 0.01; and ###p < 0.001; and ####p < 0.0001.

SYT1 is a multifunctional protein that also serves to clamp spontaneous neurotransmitter release under resting conditions (Bai et al., 2016; Chicka et al., 2008; Courtney et al., 2019; Liu et al., 2014; Vevea & Chapman, 2020). To investigate whether over-expressed pH-SYT9 can clamp spontaneous fusion, miniature inhibitory postsynaptic currents (mIPSCs, also called minis) were recorded in the presence of tetrodotoxin, to block action potentials (**Fig. 2e**). Disruption of SYT1 resulted in a ∼3-fold increase in mIPSC frequency with no rescue observed in the 1x pH-SYT9 over-expression condition. In contrast, 25x over-expression of SYT9 resulted in a complete rescue of clamping activity (**Fig. 2f**). mIPSC amplitude and rise and decay kinetics were unaffected across conditions (data not shown). Taken together, these findings indicate that SYT9 can rescue evoked release and clamp spontaneous release, in *Syt1* KO neurons, only when massively over-expressed.

To investigate how the localization of over-expressed SYT9 functions to support SV release, we utilized the N-terminal pHluorin tag to differentiate between pools in the plasma membrane versus internal compartments (see Methods and [Courtney et al., 2019]). We co-stained with antibodies against synaptophysin (SYP), to label SVs. The internal fraction in the 1x over-expression condition showed some degree of colocalization with SYP, and the internal fraction of 25x pH-SYT9 had a higher degree of colocalization. We note that endogenous SYT9 colocalized with SYP to a similar degree as 1x over-expressed pH-SYT9, while 25x over-expressed pH-SYT9 colocalized with SYP to a similar degree as endogenous SYT1 (**Fig. 2h, i**). These results are congruent with the idea that some SYT9 may be present on SVs, as indicated in proteomic studies (Takamori et al., 2006; Taoufiq et al., 2020). However, these levels are too low to support evoked release or to clamp spontaneous release, at least in the types of neurons studied here.

### Decreased mIPSC frequency in SYT9 KO striatal neurons

To further characterize neurotransmission in *Syt9* KO mice, spontaneous release from cultured striatal neurons was measured and compared to WT controls. We reiterate that all aspects of release evoked by single action potentials were normal (**Fig. 1g**); however, the mIPSC frequency in *Syt9* KO striatal neurons was reduced 2-fold (**Fig. 3a,c**), with no effect on the amplitude or kinetics of these spontaneous quantal events (**Fig. 3b**). Interestingly, when the same experiment was performed using cortical and hippocampal neurons, loss of SYT9 did not affect spontaneous release rates (**Supp. Fig. 4**), suggesting that this phenotype was specific for striatal neurons, most of which are GABAergic medium spiny neurons (MSNs) (Kemp & Powell, 1971). Importantly, the density of synapses was unchanged between WT and *Syt9* KO striatal neurons (**Supp. Fig. 5**).

**Figure 3.**
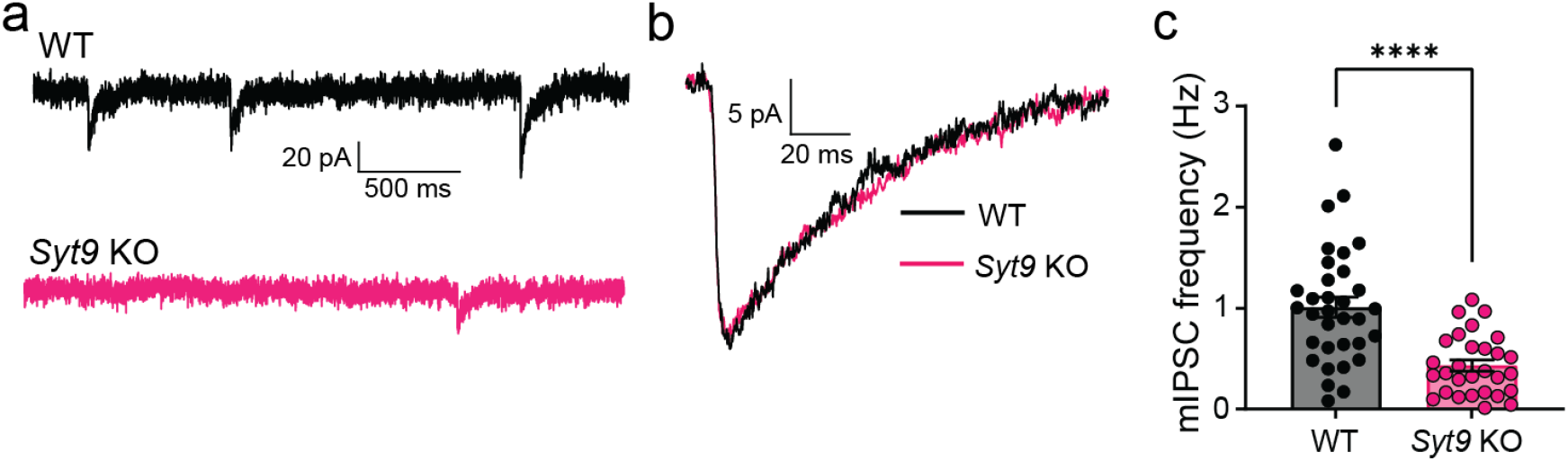
SYT9 KO striatal neurons display decreased mIPSC frequency. **a**, Representative mIPSCs traces recorded from WT (n = 33) and *Syt9* KO (n = 29) striatal neurons. **b**, Overlay of averaged mIPSCs from the two above conditions, revealing no change in kinetics or amplitude. **c**, mIPSC frequency is reduced in *Syt9* KO striatal neurons (p < 0.0001; Welch’s t-test).

We note that most mIPSCs are Ca^2+^-dependent and are thought to use the same Ca^2+^ sensors as evoked synchronous and asynchronous release, including SYT1 and Doc2 (Courtney et al., 2018; Groffen et al., 2010; Xu et al., 2009). The insufficiency of endogenous SYT9 to drive evoked release argues against the idea that it is a Ca^2+^ sensor for mIPSCs. Furthermore, the specificity of the mIPSC phenotype for striatal versus hippocampal or cortical neurons suggests a cell-specific mechanism for the action of SYT9 in spontaneous release. Notably, SYT9 is targeted to DCVs in a variety of tissues (Fukuda et al., 2002; Iezzi et al., 2005; Lynch & Martin, 2007; Roper et al., 2015), and some DCV neuropeptides have been shown to regulate aspects of synaptic transmission (Cao et al., 2011; Dean et al., 2009; He et al., 2019). This prompts the idea that SYT9 might regulate minis indirectly, potentially by controlling the release of neuromodulators from DCVs.

### SYT9 regulates the secretion of substance P-pHluorin from striatal neurons

To investigate the mechanism by which SYT9 regulates mIPSC frequency, we conducted further localization studies. Since the *Syt9* KO phenotype was specific for striatal neurons, we focused on striatum-specific DCV neuropeptides. Among these, substance P (SP) was of particular interest because it has been shown to regulate mIPSC frequency in striatal neurons through its action on the neurokinin 1 receptor (NK1R) (He et al., 2019). Interestingly, SP is not enriched in hippocampal or cortical neurons (Shults et al., 1984), where the mIPSC phenotype was not observed.

Specific antibodies for SP are not available, so we transduced striatal neurons with a lentivirus that expresses an SP-pHluorin (SP-pH) fusion protein and performed immunocytochemistry (ICC) to assess co-localization of the pHluorin tag with endogenous SYT9. Strikingly, SYT9 was colocalized with SP-pH (**Fig. 4a,b**). The correct localization of the SP-pH construct was confirmed by staining for chromogranin B (CHGB), a canonical DCV marker in neurons (**Fig. 4c,d**). In summary, these findings demonstrate the presence of SYT9 on DCVs that bear a neuropeptide known to regulate mIPSC frequency (He et al., 2019)

**Figure 4.**
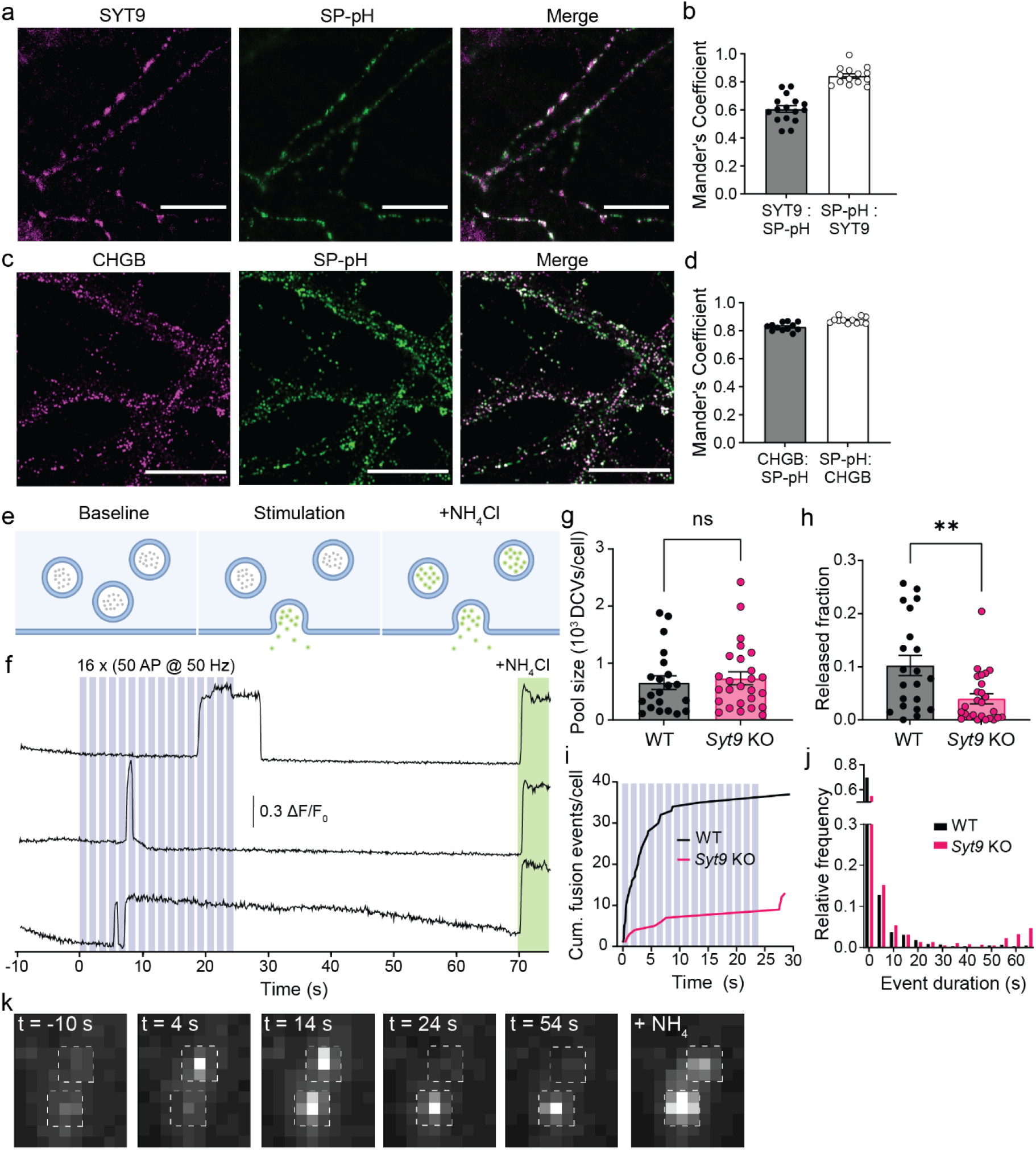
SYT9 regulates the secretion of substance P-pHluorin from striatal neurons. **a**, Representative images of cultured striatal neurons stained with α-SYT9 (magenta) and α-GFP (SP-pH) (green) antibodies; channels are merged on the right. **b**, SYT9 is co-localized with SP-pH, and vice versa; quantification was conducted using the Mander’s overlap coefficient. **c**-**d**, As in panels (**a-b**) but co-stained with α-CHGB (magenta) and α-GFP antibodies (green). SP-pH is co-localized with CHGB, confirming that SP-pH is targeted to DCVs. **e**, Schematic of the SP-pH release assay. **f**, Representative traces of individual SP-pH vesicle fusion events; the stimulation paradigm is overlaid in blue, and alkalinization of DCVs with NH_4_Cl is shown in green. **g**, Pool size per neuron was unchanged in *Syt9* KO neurons (p = 0.60; unpaired t-test). **h**, The released fraction of SP-pH is reduced in *Syt9* KO neurons (n = 27) compared to WT (n = 21) (p = 0.0049; Mann-Whitney test). **i**, Plot of cumulative SP-pH fusion events; again, the stimulus protocol is indicated in blue. **i**, Histogram of SP-pH release event duration for WT and SYT9 KO striatal neurons. **k**, Representative regions of interest (ROIs) in a timelapse, demonstrating changes in SP-pH fluorescence over the course of an imaging experiment, taken from **Video 1**; time points are indicated, with the stimulus at t = 0. The last image shows the same ROI after perfusion with NH_4_Cl to neutralize the luminal pH of SP-pH-bearing DCVs, as indicated in (**f**). **p < 0.01

Next, we sought to determine whether SYT9 regulates SP release from DCVs in striatal neurons. To optically measure SP secretion events at single-event resolution, WT and *Syt9* KO neurons were sparsely transfected with a plasmid encoding SP-pH to enable imaging of individual neurons. SP-pH fluorescence is quenched in the low pH environment of the DCV lumen; upon exocytosis, the pHluorin is rapidly dequenched, yielding a robust increase in fluorescence intensity (**Fig. 4e**). In cultured neurons, repetitive high-frequency electrical stimulation (i.e.,16 bursts of 50 APs at 50Hz) has been shown to effectively trigger these release events (Persoon et al., 2018); this paradigm was used in our experiments and yielded robust exocytosis (**Fig. 4f; Video 1**). DCV fusion was quantified in WT and *Syt9* KO striatal neurons, where the released fraction was calculated as the number of fusion events divided by the total number of SP-pH vesicles in a single neuron. The total SP-pH DCV pool was estimated by perfusing neurons with NH_4_Cl to alkalinize the lumen of DCVs and dequench the pHluorin reporter; no significant differences were observed between the WT and *Syt9* KO (**Fig 4g**). In WT neurons the released fraction was ∼10%. In *Syt9* KO neurons this was reduced by ∼61% (**Fig. 4h, i**), with only ∼3.9% of the vesicles being released. Fusion event duration was also measured in WT and *Syt9* KO neurons, and there appeared to be a small population of long-duration release events that was absent in the WT (**Fig. 4j**), hinting at a potential role of SYT9 in regulating the fate of fusion pores, as shown for other SYT isoforms (Rao et al., 2014; Wang et al., 2001). Finally, loss of SYT9 had no apparent effect on the transport of SP-pH vesicles, suggesting that the observed reductions in SP secretion were not secondary to trafficking defects (**Supp. Fig. 6**).

### Application of exogenous substance P rescues the SYT9 KO mIPSC phenotype

Since our data suggest that SYT9 regulates SP release (**Fig. 4**), and SP is known to modulate mIPSC frequency in striatal neurons (He et al., 2019), we sought to determine whether the *Syt9* KO phenotype could be rescued via acute application of this neuropeptide. In addition to SP, enkephalins, dynorphins, and cholecystokinin are also enriched in the striatum (Angulo & McEwen, 1994; Krause et al., 1987), so we also assayed the effects of each of these neuropeptides (**Fig. 5a**). Acute addition of SP to the bath fully rescued the decreased mIPSC frequency in *Syt9* KO neurons. Rescue was also observed using GR73632, a synthetic NK1R agonist. In contrast, dynorphin A (DynA), Leu-enkephalin (L-Enk), and cholecystokinin (CCK8) (**Fig. 5b**) had no effect, demonstrating that the mIPSC phenotype is SP-specific. Also, we found that SP supplementation did not significantly change the mIPSC frequency of WT neurons, indicating a high degree of SP-receptor activation under basal conditions.

**Figure 5.**
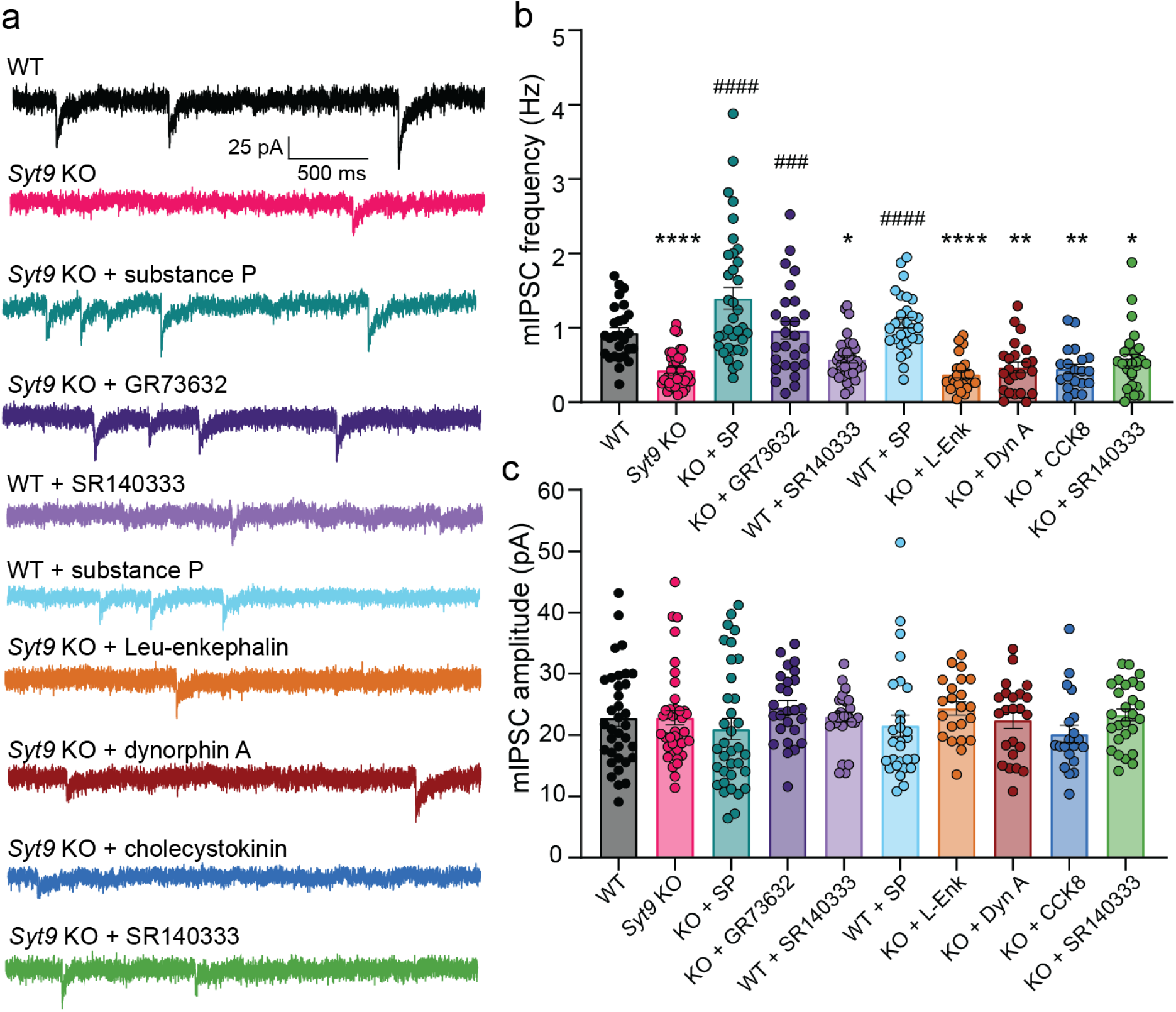
Treatment with substance P rescues SYT9 KO mIPSC phenotype. **a**, Representative mIPSC traces recorded from striatal neurons as follows: WT (n = 35); *Syt9* KO (n = 39); WT with application of 100 nM SP (n = 28) or 10 nM SR140333 (n = 34); *Syt9* KO with application of either: 100 nM SP (n = 35), 100 nM GR73632 (n = 26), 10 nM SR140333 (n=27), 100 nM Leu-enkephalin (n = 23), 100 nM dynorphin A (n = 23), or 100 nM cholecystokinin (n = 20) in the bath solution. **b**-**c**, Average amplitude (**b**) and frequency (**c**) of mIPSCs recorded from (**a**). Asterisks (*) indicate statistical comparisons (one-way ANOVA with Šídák’s multiple comparisons test) to WT neurons; hash symbols (#) indicate statistical comparisons to *Syt9* KO neurons. *p < 0.05, **p < 0.01; ****p < 0.0001; ###p < 0.001 and ####p < 0.0001.

Next, we further explored the SP-NK1R pathway using a pharmacological approach. Application of SR140333, a synthetic NK1R antagonist (Jung et al., 1994), to WT neurons, phenocopied the *Syt9* KO; mIPSC frequency was reduced ∼2-fold (**Fig. 5b**). This finding is consistent with a previous study utilizing a different NK1R antagonist (He et al., 2019), and confirms that the SP-NK1R pathway serves to regulate mini frequency in striatal neurons. Consistent with this model, SR140333 had no effect on mini frequency in *Syt9* KO neurons, presumably due to the large reduction in SP release from these cells and the concomitant decrease in NK1R activation. No differences in mIPSC amplitude were observed across all conditions (**Fig. 5c**). In summary, SYT9 regulates SP release, and activation of the NK1R by SP is pivotal in the regulation of mIPSCs in striatal neurons.

The binding of SP to the NK1 receptor presumably activates G_q/11_, resulting in the generation of IP_3_ and the mobilization of Ca^2+^ from internal stores. While basal presynaptic [Ca^2+^]_i_ did not appear to be altered in the *Syt9* KO striatal neurons (**Supp. Fig. 7**), it has been shown that local, transient increases in [Ca^2+^]_i_, mediated by IP_3_ receptors in the presynaptic ER, drive spontaneous neurotransmitter release in pyramidal neurons (Emptage et al., 2001; Simkus & Stricker, 2002). We therefore speculate that decreased SP-mediated IP_3_ signaling in the *Syt9* KO underlies the decreased mIPSC phenotype observed in our studies.

### Ca^2+^-binding to the C2A domain of SYT9 is crucial for its function *in vitro* and in neurons

The findings presented thus far suggest a model in which SYT9 regulates SP release which, in turn, modulates mini frequency. To further evaluate this model and directly determine whether SYT9 serves as a Ca^2+^ sensor for SP release, we sought to identify the structural elements in SYT9 that regulate membrane fusion *in vitro*, and then correlate these findings with the function of SYT9 in regulating both SP release and mini frequency. For these experiments, we used soluble, cytoplasmic fragments of SYT9, and for comparison, SYT1, in a ‘split t-SNARE’ *in vitro* fusion assay (Bhalla et al., 2006) that mimics fusion of SVs with the plasma membrane in cells. In this assay, SYT fragments must first fold soluble SNAP-25B onto membrane-embedded syntaxin 1A (SYX1A) for fusion with synaptobrevin 2 (SYB2)-bearing liposomes to occur. Consistent with a previous study (Roper et al., 2015), the cytoplasmic domains (C2AB) of both SYT1 and SYT9 stimulated fusion to similar extents in the presence of Ca^2+^ (omitted for clarity in **Fig. 6a** left panel but quantified in **Fig. 6a** right panel). We extended these experiments by evaluating the isolated C2-domains of each isoform and found that the C2A domain of SYT9 strongly stimulated fusion in response to Ca^2+^, while C2B had little effect (**Fig. 6a**). This is in sharp contrast to SYT1, where C2B, but not C2A, facilitated fusion (**Fig. 6a**) (Gaffaney et al., 2008). Additionally, we utilized Ca^2+^ ligand mutants (CLMs), in which two native Asp residues that serve as conserved Ca^2+^ ligands in numerous isoforms were substituted with Asn, in either C2A or C2B of SYT9. These mutations disrupted the Ca^2+^ binding activity (**Fig. 6c**) of each C2-domain and abolished the ability of the C2A domain of SYT9 to stimulate membrane fusion in response to Ca^2+^ (**Fig. 6a**). Hence, these two related isoforms have diverged unexpectedly, with C2A serving as the primary Ca^2+^ sensing domain in SYT9, while C2B is more crucial in SYT1.

**Figure 6.**
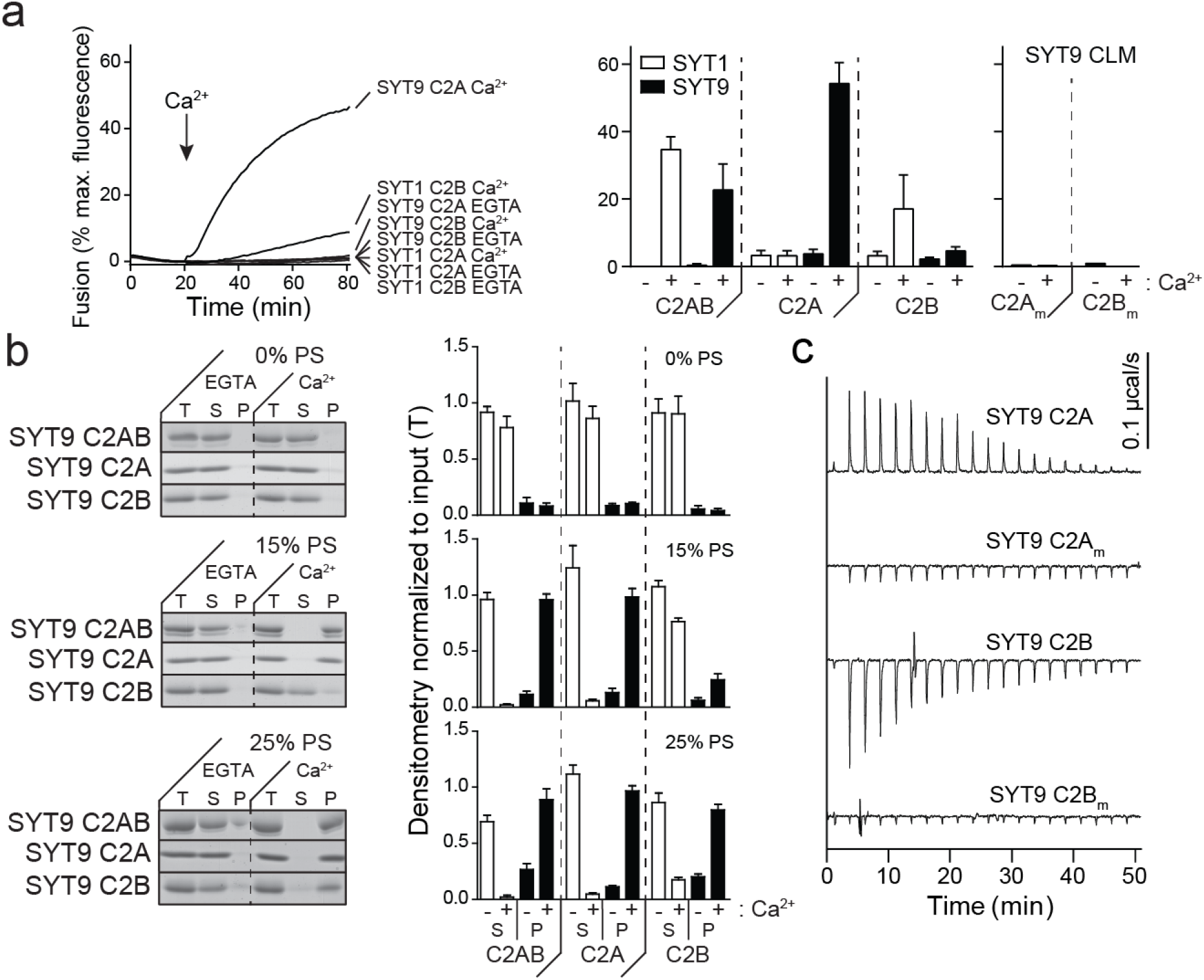
Ca^2+^•C2A from SYT9 regulates membrane fusion *in vitro*. **a**, *Left panel:* Reconstituted split t-SNARE fusion assay where Ca^2+^•SYT fragments must fold soluble SNAP-25B onto membrane-embedded SYX1A for fusion with SYB2 proteoliposomes to occur. Representative fusion traces, using each isolated C2-domain, C2A and C2B, from SYT1 and SYT9 are shown (left); the tandem C2-domains, C2AB, for each isoform were also assayed in parallel but were omitted for clarity. The isolated C2A domain, but not the C2B domain, of SYT9 stimulated fusion; in SYT1, C2B, rather than C2A, stimulated fusion (Gaffaney et al., 2008). *Right panel*: The extent of fusion (80 min), using all SYT constructs, in the presence (+) or absence (-) of Ca^2+^, is plotted (n ≥ 3). **b**, Binding of SYT9 C2A, C2B, and C2AB to PS-bearing liposomes was monitored via a co-sedimentation assay. Representative SDS-PAGE gels of protein from equal fractions of the supernatant (S), and pellet (P), as well as the total input (T), in the absence (0.2 mM EGTA) or presence of Ca^2+^ (1 mM free), using liposomes with 0, 15 and 25% PS, are shown (left). Protein band intensities were normalized to the total input and plotted (right; n ≥ 3). **c**, Representative ITC traces showing the heat of Ca^2+^ binding to the isolated C2-domains of WT SYT9; the Ca^2+^ ligand mutant (CLM) domains were analyzed in parallel, confirming that they fail to bind Ca^2+^ (n = 4). Thermodynamic values are provided in **Supplementary Table 1**.

Since SYT1 regulates fusion via interactions with membranes that harbor anionic phospholipids (Bai et al., 2016; Bhalla et al., 2005; Liu et al., 2014), we assayed the ability of SYT9 to bind phosphatidylserine (PS)-harboring membranes in the presence and absence of Ca^2+^. A previous study reported that for SYT9 C2A, but not C2B, bound to PS-bearing vesicles (Shin et al., 2004). We found that each of the isolated C2-domains efficiently bound to liposomes, in response to Ca^2+^, when the mole fraction of PS was increased from 15% to 25%; this interaction is physiologically relevant since PS is ∼22% of the inner leaflet of the plasma membrane (Hamilton et al., 2000) (**Fig. 6b**).

We also directly measured Ca^2+^ binding to the isolated C2-domains of SYT9, using isothermal titration calorimetry (ITC); calorimetry data for SYT1 has been previously reported (Evans et al., 2016; Radhakrishnan et al., 2009). Ca^2+^ binding to SYT9 C2A was endothermic and binding to C2B was exothermic (**Fig. 6c**). When fitted with a sequential binding site model, C2A exhibited three binding sites with K_d_ values of 53 µM, 182 µM, and 337 µM; C2B had two binding sites, with K_d_ values of 43 µM and 500 µM (see **Supplementary Table 1** for all thermodynamic values). Interestingly, the K_d_ value for the third binding site of C2A (337 µM) was significantly lower than the value reported for SYT1 (K_d_ > 1 mM (Evans et al., 2016; Radhakrishnan et al., 2009; Ubach et al., 1998). Titrations were also performed on the CLM forms of isolated C2A or C2B, described above, and these constructs yielded little to no signal, confirming they fail to bind Ca^2+^ (**Fig. 6c**). These experiments demonstrate that while both isolated C2-domains bind Ca^2+^, only C2A was able to effectively stimulate membrane fusion.

Next, if our model is correct, the biochemical results above suggest that Ca^2+^ binding to C2A, but not C2B, should underlie the ability of SYT9 to promote SP release. To test this, full-length variants of the SYT9 CLMs were generated and expressed in SYT9 KO neurons via lentiviral transduction; WT SYT9 served as a positive control. SP release was monitored using the optical assay described in **Figure 4**. Importantly, C2A_m_B and C2A_m_B_m_ failed to rescue release, while C2AB_m_ exhibited full rescue (**Fig. 7a**) with no change in DCV pool size (**Fig. 7b**). Hence, the ability of these mutants to regulate reconstituted membrane fusion reactions *in vitro*, and to drive SP release from living neurons, were well correlated; in both cases, the C2A domain functioned as the crucial Ca^2+^-sensing motif. These findings further support the idea that SYT9 functions as a Ca^2+^ sensor for DCV exocytosis.

**Figure 7.**
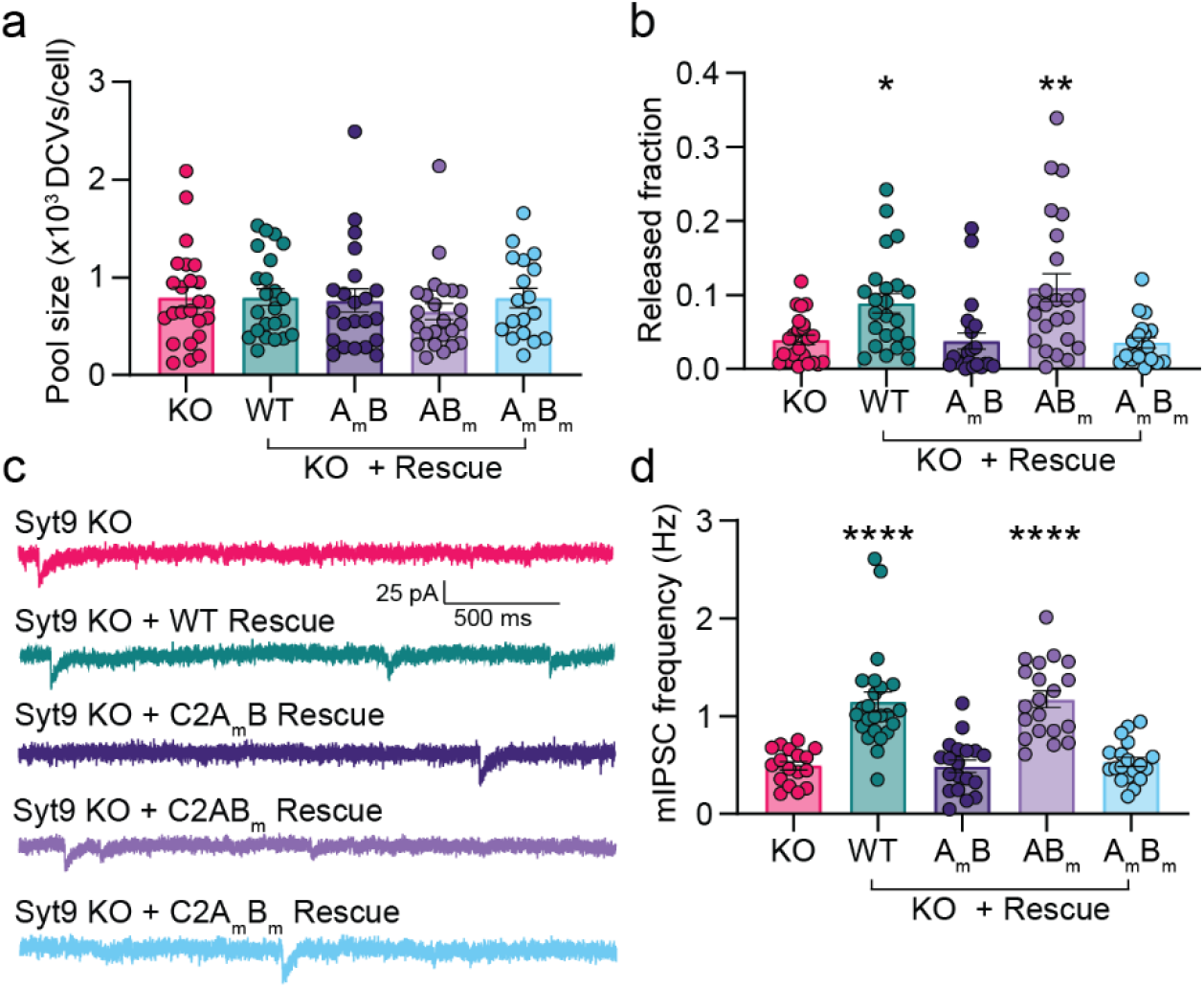
The Ca^2+^-binding activity of the C2A domain of SYT9 is required for its action in neurons. **a**, SP-containing DCV pool size was unchanged across *Syt9* KO rescue conditions (p = 0. 6272; Kruskal-Wallis test). **b**, The reduction in SP-pH release in *Syt9* KO striatal neurons (n = 21) was rescued by expression of WT SYT9 (n = 23) (p = 0.0158) or the C2AB_m_ mutant (n = 24) (p = 0.0069). In contrast, the C2A_m_B (n = 22) (p > 0.9999) and C2A_m_B_m_ mutants (n = 19) (p > 0.9999) both failed to rescue release. Release was monitored as shown in (**Fig. 4e-k**). **c**, Representative mIPSC traces recorded from striatal neurons as follows: *Syt9* KO (n = 17), *Syt9* KO + WT rescue (n = 24), *Syt9* KO + C2A_m_B rescue (n = 19), *Syt9* KO + C2AB_m_ rescue (n = 20), and *Syt9* KO + C2A_m_B_m_ (n = 19) rescue. **d**, Average frequency of mIPSCs recorded from (**c**). WT SYT9 and C2AB_m_ mutant rescued the mini frequency, while the C2A_m_ and C2A_m_B_m_ mutants failed to rescue this phenotype. In this figure, asterisks (*) indicate statistical comparisons (Kruskal-Wallis test with Dunn’s multiple comparisons test) to the *Syt9* KO condition.

Finally, we conducted patch-clamp recordings of *Syt9* KO striatal neurons that expressed each full-length CLM construct. Again, complete rescue of mIPSC frequency was observed using WT SYT9. In sharp contrast, SYT9 C2A_m_B_m_, which cannot bind Ca^2+^ via either C2-domain, failed to rescue the mini phenotype. Consistent with our model, C2A_m_B also failed to rescue (**Fig. 7c**), while C2AB_m_ was as effective as the WT protein (**Fig. 7d**; note: no changes in mini amplitude were observed across conditions [**Supp. Fig. 8**]). In summary, Ca^2+^ binding to the C2A domain, but not the C2B domain, underlies the ability of SYT9 to regulate membrane fusion to, in turn, indirectly control the rate of spontaneous release in striatal neurons by controlling SP secretion.

## DISCUSSION

SYT9 is widely expressed in the brain ([Mittelsteadt et al., 2009]; note: these authors referred to SYT9 as SYT5), but KOs lacking this protein have only been characterized using the calyx of Held slice preparation (Kochubey et al., 2016) and cultured striatal neurons (Xu et al., 2007). SYT9 appears to be expressed in the calyx of Held, but loss of this protein did not affect AP-triggered neurotransmitter release (Kochubey et al., 2016). In contrast, in cultured striatal neurons, Xu et al. (2007) reported a 57% reduction in eIPSC amplitude in *Syt9* KO striatal neurons. However, this finding has not been reproduced, and it is difficult to reconcile with the expression patterns of SYT9 and SYT1. Namely, SYT9 is expressed at similar levels in cortical, hippocampal, and striatal neurons, and it is well-established that loss of SYT1 abolishes synchronous release in both cortical and hippocampal neurons (Geppert et al., 1994; Maximov & Südhof, 2005) (**Fig 1**). So, it is apparent that endogenous SYT9 does not function as a Ca^2+^ sensor for synchronous SV exocytosis in either of these two classes of neurons (**Fig. 1; Supp. Fig. 1**). Here, we extended these observations to cultured striatal neurons and found that SYT1, but not endogenous SYT9, drives rapid SV exocytosis. We went on to show that SYT9 is largely localized to SP-bearing DCVs in striatal neurons, where it regulates the release of this neuropeptide to indirectly control mIPSC frequency. Evidently, at least some isoforms of SYT have become specialized to control the release of neuropeptides that impact synaptic function (Cao et al., 2011; Dean et al., 2009; Dean, Liu, et al., 2012). It is tempting to speculate the divergence of numerous SYT isoforms might enable the differential release of myriad distinct neuromodulators and hormones, in response to different patterns of activity.

We found that SYT9 is expressed at ∼25-fold lower levels than SYT1 (**Fig. 1**). Given the estimate of ∼15 copies of SYT1 per SV (Takamori et al., 2006), the abundance of SYT9 on SVs would, on average, be expected to be low, particularly since its primary localization appears to be on DCVs (**Fig. 4**). Some fraction of SYT9 does colocalize with canonical SV markers (**Fig. 2**; (Xu et al., 2007), and SYT9 was detected in proteomic studies of purified SVs ([Takamori et al., 2006; Taoufiq et al., 2020] note: these authors referred to SYT9 as SYT5). Thus, it remains possible that there could be limited populations of SVs, in discrete brain regions, which harbor a sufficient copy number of SYT9 molecules to trigger SV exocytosis. However, in our hands, this was not the case with cortical, hippocampal, or striatal neurons (**Fig 1; Supp. Fig. 1**). It should also be emphasized that DCVs are often present in or near presynaptic nerve terminals (Persoon et al., 2018; Radhakrishnan et al., 2021), and this can potentially exaggerate the presence of SYT9 on SVs as determined via immunocytochemistry.

While our experiments indicate that SYT9 functions as a Ca^2+^ sensor for SP release from striatal neurons, some degree of regulated release persisted in the KO (**Fig. 4**), suggesting that additional Ca^2+^ sensors are associated with SP-bearing DCVs. Indeed, SYT1 was present on SP-bearing DCVs purified from rabbit optic nerve (Berg et al., 2000). Additionally, in hippocampal neurons, SYT1 and SYT7 have both been implicated in DCV exocytosis (van Westen et al., 2021), suggesting multiple roles for these Ca^2+^ sensors. We also note that SYT4, which does not bind Ca^2+^, negatively regulates the release of brain-derived neurotrophic factor (BDNF) (Dean et al., 2009), and SYT10, which binds Ca^2+^, positively regulates the secretion of IGF-1 from DCVs in neurons (Cao et al., 2011). Collectively, these findings, combined with the observations in the current study, support the idea that the SYTs have diverged, at least in part, to control different forms of DCV exocytosis (Dean, Dunning, et al., 2012). We also reiterate that the *Syt9* KO mIPSC phenotype appears to be specific for striatal neurons, which robustly express SP. The function of SYT9 in cortical and hippocampal neurons remains unknown. We speculate that it regulates the release of neuropeptides; a future goal is to address this issue via co-localization studies and by isolating and characterizing SYT9-bearing organelles from these classes of neurons.

Regarding the diversity of SYT isoforms, our finding that the C2A domain of SYT9 serves as the crucial Ca^2+^-sensing motif for fusion was unexpected. SYT9 forms a clade with SYT1/2, and Ca^2+^ ligand mutations in the C2B-domain of SYT1 clearly disrupt function (Nishiki & Augustine, 2004), while similar mutations in C2A have been reported to either have no effect (Fernández-Chacón et al., 2002), to be gain-of-function (Pang, Shin, et al., 2006; Stevens & Sullivan, 2003), or to cause partial loss-of-function (Shin et al., 2009). Moreover, the isolated C2B domain of SYT1, but not the C2A domain, regulates fusion *in vitro* (**Fig. 6**; (Gaffaney et al., 2008). In contrast, Ca^2+^-binding to the C2A domain of SYT9, but not the C2B domain, was required to both stimulate fusion *in vitro* (**Fig. 6**) and promote SP release from striatal neurons (**Fig. 7a,b**), to regulate minis (**Fig. 7c,d**). Apparently, this shuffling of C2-domain function is tolerated, as SYT9 can rescue evoked neurotransmitter release and clamp spontaneous release in *Syt1* KO neurons, when sufficiently over-expressed (**Fig. 2**).

Returning to the physiology, we emphasize that the striatum is a specialized structure with a dense local network largely composed of medium spiny neurons (MSNs) and their axon collaterals. While a substantial fraction of these neurons projects to other brain regions, direct synaptic communication between MSNs also occurs (Koos et al., 2004; Tunstall et al., 2002); this is the condition recapitulated in our experiments using cultured striatal neurons. Subpopulations of MSNs release SP within the striatum (Gerfen, 1992) and harbor NK1 receptors (Almeida et al., 2004; Haneda et al., 2007). We observed modulation of MSN-MSN transmission by SP in culture, and the next step will be to measure such modulation in acute striatal slice preparations. In addition, MSNs project to targets in the substantia nigra and globus pallidus, and recordings from brain slices that encompass the striatum along with these other structures will address the role of SYT9-regulated SP release at terminals that innervate these target cells. Moreover, SP enhances center-surround contrast of striosomal dopamine signals by modulating dopamine release from neurons that project from the substantial nigra onto striatal targets (Brimblecombe & Cragg, 2015). Hence, slice recordings, to measure dopamine release, are an important future direction of this work.

We propose that SP modulates spontaneous release rates in striatal neurons by binding the NK1 receptor and activating G_q/11_, resulting in the generation of IP_3_ and the mobilization of Ca^2+^ from internal stores (**Fig. 8**), thus increasing mini frequency. Indeed, localized increases in [Ca^2+^]_i_, mediated by IP_3_ receptors in the presynaptic ER, have been reported to drive spontaneous release in pyramidal neurons (Emptage et al., 2001; Simkus & Stricker, 2002). A future goal is to compare these brief Ca^2+^ transients, akin to Ca^2+^ sparks in muscle cells, in WT and *Syt9* KO striatal neurons. Finally, we emphasize that the highest expression levels of SYT9 in mouse brain occur in the pituitary gland and hypothalamus, both of which are particularly rich sources of numerous classes of DCVs (Roper et al., 2015). It will be interesting to investigate whether SYT9 regulates the release of hypothalamic hormones.

**Figure 8.**
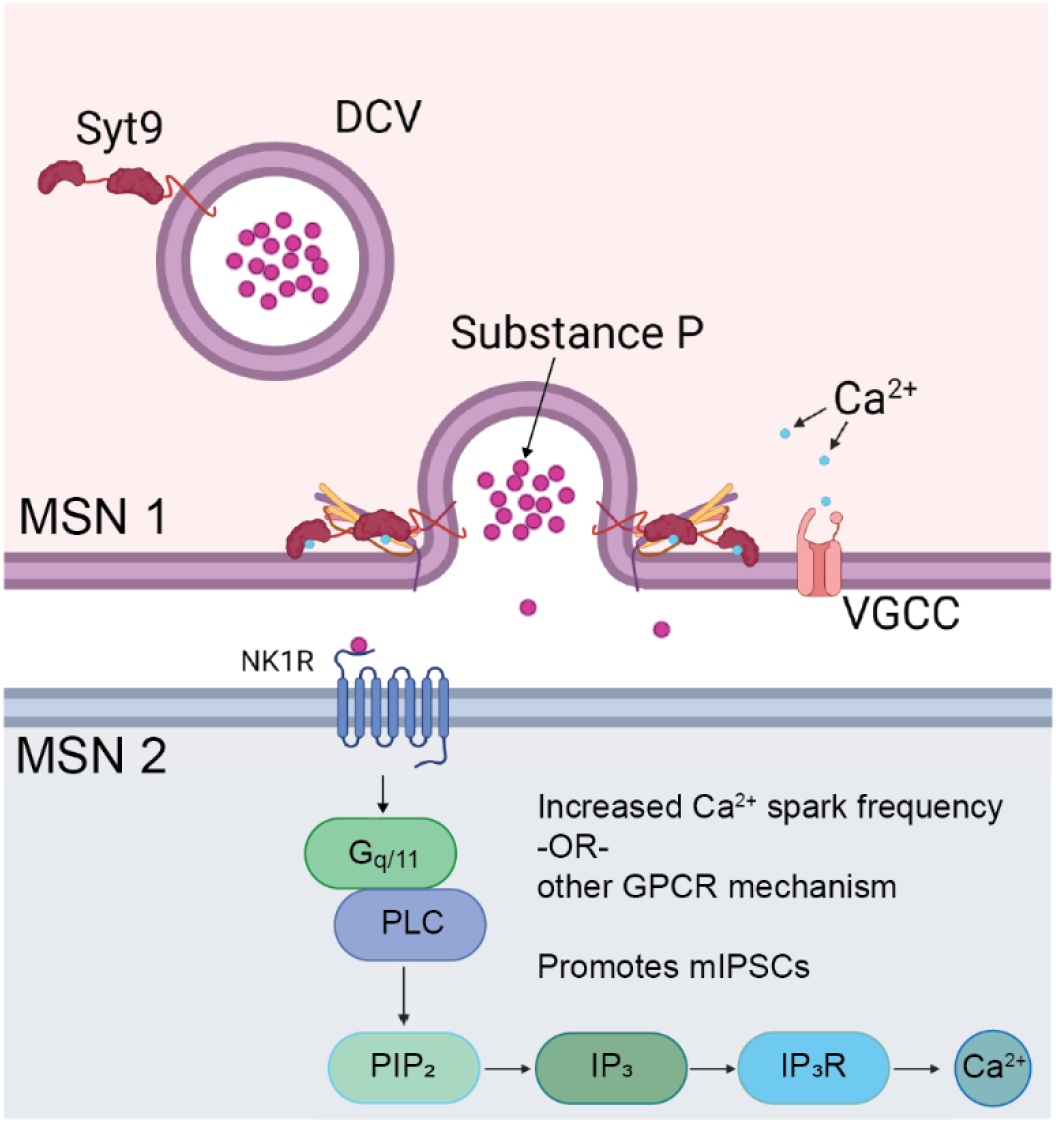
SYT9 indirectly controls mini frequency in striatal neurons by serving as a Ca^2+^ sensor for SP exocytosis. A model in which SYT9 serves as a Ca^2+^ sensor that regulates the exocytosis of SP-containing DCVs from striatal neurons. Released SP then activates the NK1 receptor which in turn activates the IP_3_ pathway via G_q/11_ and phospholipase C (PLC), leading to increased Ca^2+^ efflux from the ER, to promote spontaneous neurotransmitter release rates.

## MATERIALS AND METHODS

### Animals

All neuronal cultures were prepared from early postnatal (P0-P1) mice in accordance with all relevant regulations and the approval of the University of Wisconsin Institutional Animal Care and Use Committee. Conditional *Syt9 KO* mice were obtained from Jackson Labs (Xu et al. 2007). These conditional *Syt9* KO mice were terminally crossed with an E2a-CRE mouse (Jackson Labs, Bar Harbor, ME, USA) to yield a population of constitutive *Syt9* KO mice; the mice used in this study are all products of this constitutive *Syt9* KO mouse. *Syt1* cKO (conditional KO) mice were a gift from W. Thoreson, (University of Nebraska-Lincoln, Lincoln, NE, USA; [Quadros et al., 2017]).

### Genotyping

The *Syt9* KO mouse line was maintained as heterozygous breeders so that offspring WT and KO littermates could be studied. To genotype the neonatal pups, tissue samples were taken by collecting ∼2 mm of the end of the tail and extracting DNA from the tissue sample using the Kapa Express Extract Kit (Kapa Biosystems, Wilmington, MA, USA). The extracted DNA was then subjected to PCR using in-house designed primers to identify WT and KO alleles. The primers used to identify the WT allele yield a 209 bp amplicon: F-primer 5’- GCGTCAATGGGAGGAGAGGTCA-3’; R-primer 5’-GTCCACTAGGGCTAGCCCAGG-3’. The primers used to identify the KO allele yield a 301 bp amplicon: F-primer 5’- CAGTAGGGAGCGAGCAGAAATTAAAAT-3’; R-primer 5’- AACTTCGTCGATCGACCTCGAATAA-3’. The PCR product was subjected to agarose gel electrophoresis and imaged on a ChemiDoc MP imaging system (BioRad, Hercules, CA, USA).

### Molecular biology and lentiviral preparation

Lentiviral vectors were based on FUGW (FUGW was a gift from David Baltimore [Addgene plasmid # 14883; http://n2t.net/addgene:14883; RRID: Addgene_14883]) (Lois et al., 2002). The FUGW vector was modified by substituting the ubiquitin promoter with the human synapsin I promoter (Kügler et al., 2003); this vector is henceforth referred to as FSGW. pH-SYT9 was generated by replacing SYT1 in pHluorin-SYT1 (a gift from J. Rothman, Yale University, New Haven, CT) (Fernández-Alfonso et al., 2006) with a SYT9 cDNA (a gift from M. Fukuda, Tohoku University, Sendai, Miyagi, Japan) (Haberman et al., 2003) using PCR splicing via overlap extension (SOE). The pHluorin-SYT9 cassette replaced GFP in the cloning site of the FSGW vector via In-Fusion Cloning (Takara Bio, San Jose, CA, USA). The SP-pH construct was generated by PCR SOE and comprised: the first 68 residues of mouse preprotachykinin (PPT), a GS(GSS)_4_ linker, and a super-ecliptic pHluorin (Miesenböck et al., 1998). This fusion construct was subcloned into FSGW.

SYT9 CLM constructs correspond to D-to-N substitutions of two Ca^2+^ ligands in each C2-domain as follows: C2A_m_B; D197N and D199N; C2AB_m_; D330N and D332N. All four mutations are present in the C2A_m_B_m_ construct. These constructs were made by performing site-directed mutagenesis on the *Syt9* cDNA with the QuikChange Lightning Site-Directed Mutagenesis Kit (Agilent, Santa Clara, CA, USA). Full-length CLM constructs were used for the rescue experiments in Fig. 7, and isolated purified C2-domain CLM constructs were used for Fig. 6. cDNA for SNAP-25B was provided by M. C. Wilson (University of New Mexico School of Medicine, Albuquerque, NM, USA), and full-length synaptobrevin 2 (SYB2) and syntaxin 1A (SYX) were provided by J. Rothman (Yale University, New Haven, CT); these SNARE proteins were subcloned into pTrcHis vectors to yield His_6_-tagged proteins (Evans et al., 2015).

### Cell culture, transient transfection, and lentiviral transduction

Cortical, hippocampal, and striatal regions were obtained by microdissection and kept in ice-cold Hibernate-A (BrainBits, Springfield, IL, USA) with 2% B-27 until completion of dissection. Neural tissue was then incubated with 0.25% trypsin-EDTA (Corning Inc., Corning, NY, USA) for 25 minutes at 37°C. The tissue was washed with DMEM (Gibco, Carlsbad, CA, USA) supplemented with FBS (Atlanta Biologicals, Flowery Branch, GA, USA) and penicillin-streptomycin (Thermo Scientific, Waltham, MA, USA) and triturated 10-15 times with a P1000 pipette tip until complete dissociation of the tissue. Cells were plated at a density of 100,000 cells/cm^2^ onto poly D-lysine-coated glass coverslips. Supplemented DMEM was replaced after 1 hour with Neurobasal-A (Gibco) supplemented with B-27 (Thermo Scientific) and GlutaMAX (Thermo Scientific). For SP-pH release experiments, sparse transfection of SP-pH and a cytosolic mRuby marker was required for identifying and imaging single neurons. This was achieved using Lipofectamine LTX with PLUS reagent (Invitrogen, Carlsbad, CA, USA) on DIV 8-9. For localization studies, rescue experiments, and Ca^2+^ imaging, full coverage was desirable and was achieved via lentiviral transduction. Lentivirus encoding CRE-recombinase was added to *Syt1* cKO neurons at DIV 0-2, and lentiviruses that expressed SYT9 rescue constructs, the Ca^2+^ imaging construct synaptophysin-HaloTag (SYP-Halo) (Bradberry & Chapman, 2022), or SP-pH for localization studies, were added to neurons at DIV 5-7.

### Electrophysiological studies

Whole-cell voltage-clamp recordings of cultured neurons (DIV 14-18) were performed at room temperature in artificial cerebrospinal fluid (ACSF) containing (in mM): 128 NaCl, 5 KCl, 1.5 CaCl_2_, 1 MgCl_2_, 30 glucose, 25 HEPES, pH 7.4 with an internal solution containing (in mM): 130 KCl, 
10 HEPES, pH 7.4, 1 EGTA, 2 ATP, 0.3 GTP, 5 phosphocreatine. Recordings were performed using a MultiClamp 700B amplifier and Digidata 1550B digitizer (Molecular Devices, San Jose, CA, USA) under the control of Clampex 10 software (Molecular Devices). GABA_A_-mediated currents were pharmacologically isolated using 20 μM CNQX (Tocris Bioscience, Bristol, UK) and 50 μM D-AP5 (Tocris) in the bath solution; for mIPSC recordings, 1 μM tetrodotoxin (Tocris) was included in the bath. 5 mM QX 314 chloride (Tocris) was included in the pipette solutions for all recordings. Neurons were held at −70 mV in all experiments without correction for liquid junction potentials. Recordings were discarded if series resistance rose above 15 MΩ. For evoked recordings, a concentric bipolar electrode (FHC, Bowdoin, ME, USA) was placed 100-200 μm away from the patched soma and stimulation currents were adjusted to evoke maximal responses. For mIPSC recordings, 300 s of data were recorded for each striatal neuron, and 180 s of data were acquired for cortical and hippocampal neurons. mIPSCs were quantified for each recording using a template-matching algorithm in Clampfit (Molecular Devices). For pharmacology rescue experiments, SP (Tocris), GR73632 (Tocris), Leu-enkephalin (Tocris), dynorphin A (Tocris), and cholecystokinin (Tocris) were added to the ACSF at 100 nM, except for SR140333 (Tocris), which was added at 10 nM. Neurons were treated with peptides or drugs for ten minutes before patch-clamp experiments were conducted.

### Immunoblotting

Coverslips of cultured neurons (DIV 14-18) were rinsed 1-2 times in PBS and lysed by repeated pipetting of lysis buffer (50 mM Tris pH 8.0, 150 mM NaCl, 2% SDS, 0.1% Triton X-100, 10 mM EDTA) containing a protease inhibitor cocktail (cOmplete mini EDTA-free, Roche). For quantitative Western blots of lysates, protein concentration was determined using a bicinchoninic acid (BCA) assay (Thermo Scientific) to ensure equal amounts of protein were subjected to SDS-PAGE. Samples were then combined with 4x Laemmli sample buffer, heated to 50°C for 15 minutes, and stored at −20°C until use. For each immunoblot experiment, the samples were subjected to SDS-PAGE on 13% polyacrylamide gels containing 1% trichloroethanol (TCE). After electrophoresis, the TCE was activated by exposure to UV light (300 nm) for 45 seconds, and labeled proteins were detected via 300 nm illumination; these signals served as a protein loading control (Ladner et al., 2004). Proteins were transferred to a PVDF membrane, which was then blocked with 5% nonfat milk in Tris-buffered saline plus 0.1% Tween 20 (TBST) for 1 hour, followed by incubation with primary antibody overnight at 4°C. Blots were then washed 3 times for 5 minutes in TBST, incubated with secondary antibody for 1 hour, and washed again as above. For the pH-SYT9 rescue experiments, after blotting for SYT9, the PVDF membranes were stripped using Restore™ Western Blot Stripping Buffer (Thermo Scientific), and the membranes were once again blocked and subsequently probed for SYT1. Primary antibodies were: α-SYT1 (1:500) (DSHB; #mAB 48; RRID:AB_2199314) and α-SYT9 (1:500) (Synaptic Systems Cat# 105 053, RRID:AB_2199639). Secondary antibodies were goat α-mouse IgG-HRP (Biorad, 1706516; RRID:AB_11125547), goat α-mouse IgG2b-HRP (Invitrogen, M32407; RRID:AB_2536647), and goat α-rabbit IgG-HRP (Biorad, 1706515, RRID:AB_11125142). Immunoreactive bands were visualized using SuperSignal West Pico Plus Chemiluminescent Substrate (Thermo Scientific) and imaged with an Amersham Imager 680 imaging system (GE Healthcare, Chicago, IL, USA). Bands were analyzed by densitometry, with care to stay within the linear range, and contrast was linearly adjusted for publication using ImageJ (Fiji) (Schindelin et al., 2012).

### Immunocytochemistry and confocal microscopy

Immunocytochemistry was conducted as previously described (Courtney et al., 2019). For the pH-SYT9 rescue experiments, immediately before fixation and permeabilization, cortical neurons were incubated in primary antibody directed against the pHluorin tag of pH-SYT9 (1:250) (α-GFP; Abcam Cat# ab290, RRID: AB_303395) in culture medium to label only the surface-resident pH-SYT9, as the antibody is unable to enter intact, live neurons. Unbound antibody was removed by washing, and then cells were fixed and permeabilized and incubated a second time with primary antibodies to label synaptophysin (1:1000) (α-SYP; Synaptic Systems Cat# 101 004; RRID:AB_1210382) and pHluorin (1:1000) (α-GFP; Abcam Cat# ab13970, RRID:AB_300798). This second α-GFP antibody labeled all pHluorin tags that were not labeled by the first α-GFP ‘blocking’ antibody, to reveal the internal fraction of pH-SYT9. This strategy allowed for differential secondary labeling of the different pH-SYT9 populations with species-specific secondary antibodies.

Images were acquired on a Zeiss LSM 880 laser scanning confocal microscope with a 63x Plan-Apochromat 1.4NA DIC oil immersion objective. Identical laser and gain settings were used for all samples within each experimental paradigm. Primary antibodies were: α-GFP (1:1000) (Abcam Cat# ab290, RRID:AB_303395), α-GFP (1:1000) (Abcam Cat# ab13970, RRID:AB_300798), α-SYT1 (1:500) (DSHB; #mAB 48; RRID:AB_2199314), α-SYP (1:1000) (Synaptic Systems Cat# 101 004; RRID:AB_1210382), α-SYT9 (1:500) (Synaptic Systems Cat# 105 053, RRID:AB_2199639), α-CHGB (Synaptic Systems Cat# 259 103; RRID:AB_2619973), α-VGAT (1:1000)(Synaptic Systems Cat # 131 004; RRID:AB_887873), α-Gephyrin (1:1000)(OriGene Cat# TA502313, RRID:AB_11126039), and α-MAP2 (1:500) (Abcam Cat# ab5392, RRID:AB_2138153). Secondary antibodies were: goat anti-chicken IgY-Alexa Fluor 405 (1:500) (abcam; ab175675, RRID:AB_2810980), goat α-chicken IgY-Alexa Fluor 488 (1:500) (Thermofisher A-11039, RRID: AB_2534096), goat α-guinea pig IgG-Alexa Fluor 488 (1:500) (Thermofisher; A-11073; RRID:AB_2534117), goat α-rabbit IgG-Alexa Fluor 546 (1:500) (Thermofisher; A-11010, RRID:AB_2534077), goat α-guinea pig IgG-Alexa Fluor 647 (1:500) (Thermofisher; A-11074, RRID:AB_2534118), and goat α-mouse IgG2b-Alexa Fluor 647 (1:500) (Thermofisher; A-21242; RRID:AB_2535811).

### Live cell imaging

DIV 14-18 striatal neurons were washed twice in ASCF before being mounted in an imaging chamber with parallel platinum electrodes (Warner Instruments, Hamden, CT, USA) with 500 μL of ACSF. Imaging was carried out with a 40x, 1.4 NA oil-immersion objective on an inverted epifluorescence microscope (IX81, Olympus, Tokyo, Japan) equipped with a CMOS camera (Orca Flash 4.0 V2, Hamamatsu Photonics, Hamamatsu, Japan), motorized stage (Mad City Labs, Madison, WI, USA), and a custom illumination source containing three LEDs (470, 530, 625 nm) (Thorlabs, Newton, NJ, USA). The system was controlled using Micro-Manager (Edelsteinet al. 2010). Stimulation was delivered with a stimulation box (SD9, Grass Instrument Co., West Warwick, RI, USA) controlled via a HEKA EPC10 DAQ-amplifier and PatchMaster software (HEKA, Holliston, MA, USA), which was also used to synchronize the start of image sequence acquisition. Images were collected using 2 x 2 binning (0.365 μm x 0.365 μm pixel size).

For SP-pH release imaging, an exposure time of 100 ms was used. Individual transfected neurons were identified for imaging by first detecting the cytosolic mRuby signal with the 530 nm LED and standard RFP filter set (49004, Chroma, Bellows Falls, VT, USA). SP-pH release was imaged with the 470 nm LED, a standard green fluorescent protein (GFP) filter set (49002, Chroma). Ten seconds of baseline were obtained before stimulation was given 16 periods of 50 APs at 50 Hz with 0.5 sec between each train. Seventy seconds after the start of stimulation, 500 μL of 2x NH_4_Cl ACSF, containing (in mM): 100 NH_4_Cl, 28 NaCl, 5 KCl, 1.5 CaCl_2_, 1 MgCl_2_, 30 glucose, 25 HEPES, pH 7.4, was added to dequench pHluorin fluorescence to visualize the total number of SP-pH-bearing vesicles.

For Ca^2+^-imaging, the 625 nm LED, custom three-band pass dichroic mirror (Chroma) and far-red emission filter, and a 10 ms exposure time were used. DIV 14-18 striatal neurons that were transduced with a virus that induced expression of SYP-Halo were first washed twice in ACSF before being incubated in ACSF containing 100 nM of the acetoxymethyl ester of JF_646_-BAPTA bearing a HaloTag ligand (JF_646_-BAPTA-AM-HTL)(Deo et al., 2019), a gift from L. Lavis (Janelia Research Campus, Ashburn, VA, USA). After a 20-minute incubation at 37°C, the cells were again washed with dye-free ACSF and incubated for 20 minutes at 37°C before being mounted in the imaging chamber. Five fields of view (FOV) were selected per coverslip. Three seconds of baseline were recorded before a saturating stimulus (50 AP @ 50Hz) was delivered to establish a maximum fluorescence (F_max_) for each field of view. At least five minutes elapsed between imaging each FOV to give the neurons time to recover and return to basal conditions.

### Image analysis

For colocalization analysis, the Coloc2 plugin in Fiji (Schindelin et al., 2012)was used to determine the Mander’s overlap coefficients and Pearson’s correlation coefficients; Costes’ automatic threshold regression was used for both determinations.

For synapse counting analysis, images were exported to SynD (Schmitz et al., 2011), where the image of MAP2 staining was used to create a neurite mask, then synaptic density was quantified by counting the number of overlapping VGAT and Gephyrin puncta within the neurite mask and dividing the total number by the length of dendrite measured to yield number of synapses per micron.

For SP-pH release analysis, Fiji with the DCV_pHluorin toolset (https://github.com/alemoro/DCVpHluorin_toolseet) (Moro et al., 2021) was used; 3 pixels x 3 pixels regions of interest (ROIs) were placed over each observed increase in fluorescence, and somatic events were excluded. Fusion events were defined as a sudden increase of 2 SD above the average fluorescence of the first 100 frames in the ROI (F_0_). Time lapses were also manually scored along with the DCV_pHluorin analysis algorithm to ensure a thorough and complete characterization of each release event. Pool size estimations were calculated using the same rules used for fusion event detection, after perfusion with NH_4_Cl. Analysis was blinded, and the number of fusion events, SP-pH vesicle pool size estimation, event duration, and fusion onset timing were all determined using DCV_pHluorin. Output data were processed in Excel and plotted using GraphPad Prism.

Ca^2+^-imaging analysis was performed as previously described (Bradberry & Chapman, 2022) with a modified ImageJ script taken from (Vevea & Chapman, 2020). Briefly, SYP-HaloTag-JF_646_-BAPTA traces were background subtracted, and ROIs were created based on a custom workflow to identify changes in fluorescence of the selected image series. An average intensity projection of pre-stimulus baseline frames was subtracted from a maximum intensity projection of the entire timelapse. This result was duplicated and mean filtered using a rolling ball radius of 10 pixels. The mean filtered image was subtracted from the original result, and this result was used to threshold changes in fluorescence. After thresholding, images were made binary, and a watershed function was run to separate overlapping objects. Using this procedure, objects (>10 pixels) were created and defined as ROIs. These ROIs were used to measure fluorescence changes over time from the original image series. The fluorescence traces of each ROI were then imported into AxoGraph where the basal, pre-stimulation fluorescence of each ROI (F_r_) was subtracted from the max fluorescence at indicator saturation (F_max_) to give a ΔF. The following equation was used to calculate presynaptic basal [Ca^2+^]_i_:

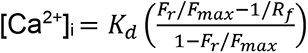

where K_d_ is the dissociation constant of the indicator, F_r_ is the baseline fluorescence of an ROI pre-stimulation, F_max_ is the fluorescence achieved with the delivery of a saturating stimulus, R_f_ is the dynamic range of the indicator, and n is the Hill coefficient. For K_d_, R_f_ and n, we used *in vitro* measurements (K_d_ = 140 nM, R_f_ = 5.5, n = 1) from (Deo et al., 2019).

Trafficking analysis was conducted using the Multi Kymograph plugin in Fiji; 30-second time-lapses, taken after NH_4_Cl perfusion in the SP-pH release experiments, were used. Ninety µm segments that began within 20 µm of the soma were chosen. The kymograph was created from this time-lapse and every path was traced and quantified.

### Protein Purification

Recombinant proteins were made as described (Evans et al., 2015). SYB2, SNAP-25B, and SYX were purified as His_6_-tagged proteins using the pTrcHis vector. The SYT1 cytoplasmic domain (SYT1 C2AB – residues 96-421; the amino acid residues of the other SYT constructs are indicated in parentheses), SYT1 C2A (96-264), and SYT1 C2B (248-421), were purified as GST-tagged proteins using a pGEX vector. The SYT9 cytoplasmic domain (SYT9 C2AB –104-386), SYT9 C2A (124-211), and SYT9 C2B (253-344) were purified as His_6_-tagged proteins using the pTrcHis vector. Isolated SYT9 C2A and C2B domains, harboring Ca^2+^ ligand mutations, were purified in the same manner at the WT C2-domains. Full-length SYT9 was purified as a His_6_-SUMO-tagged protein using the pET28 vector.

### Vesicle Preparation

Protein free liposomes, for co-sedimentation assays, were generated by drying lipids (30% PE and 70% PC; 15% PS, 30% PE, and 55% PC; and 25% PS, 30% PE, and 45% PC) in chloroform, under a stream of nitrogen. The film was rehydrated and suspended by vortexing in 50 mM HEPES-NaOH, 100 mM NaCl, pH 7.4, and extruded through 50 nm polycarbonate membranes to generate liposomes (Avanti Polar Lipids, Birmingham, AL, USA). SNARE-bearing vesicles, for *in vitro* fusion assays, were prepared as previously described (Bhalla et al., 2006). Lipid compositions were as follows: 15% PS, 27% PE, 55% PC, 1.5% NBD-PE, and 1.5% Rho-PE for SYB2 vesicles, and 25% PS, 30% PE, and 45% PC for SYX1A vesicles.

### *In vitro* Fusion Assays

Fusion assays between SYB2 and SYX1A vesicles were conducted using a scaled-down version of the referenced method (Bhalla et al., 2006). For split t-SNARE fusion assays, 1 μM of the cytoplasmic domain (C2AB) or 2 µM of isolated C2-domains, SYB2 vesicles, SYX1A vesicles, 0.2 mM EGTA, and 7 μM soluble SNAP-25B were incubated for 20 min at 37°C. Ca^2+^ (1 mM final free concentration) was added and the reaction was monitored for an additional 60 min. N-dodecyl-β-d-maltoside was added at the end of the experiment, to yield the maximum fluorescence signal. NBD donor fluorescence was monitored using a Synergy HT multi-detection microplate reader (Bio-Tek, Santa Clara, CA, USA). Traces were normalized to the first time point, and the maximum fluorescence signal, to determine the %F_max_.

### Co-sedimentation Assays

WT SYT9 C2A, C2B, or C2AB (4 μM) were combined with protein-free liposomes (1 mM total lipid; %PS as indicated in Fig. 6B) in 0.2 mM EGTA or 1 mM Ca^2+^ and incubated for 15 min at RT. Samples were then centrifuged in an Optima MAX-E tabletop ultracentrifuge (Beckman Coulter, Brea, CA, USA) at 184,000 x g (65,000 rpm) for 45 min at 4°C. After centrifugation, the supernatant (S), and the pellet (P) were collected, and equal fractions were subjected to SDS-PAGE and Coomassie blue staining. Bands were quantified by densitometry and plotted.

### Isothermal Titration Calorimetry (ITC)

WT and CLM forms of the isolated C2-domains of SYT9 were dialyzed overnight in ITC buffer (50 mM HEPES-NaOH, pH 7.4, 200 mM NaCl, and 10% glycerol) in the presence of Chelex-100 resin (BioRad), to remove residual divalent cations. ITC buffer was subsequently filtered and used to make Ca^2+^ and protein dilutions. Heat of binding was measured, using a MicroCal® iTC200 (Malvern Panalytical, Malvern, UK), at 25 °C, in response to twenty injections of Ca^2+^ into the sample cell containing 100 μM protein. Heat of dilution was corrected by subtracting the signal that arises when Ca^2+^ is diluted into buffer. Data were analyzed using the “sequential binding site” model in the data software package.

## Supporting information

Source data and statistics

## ACKNOWLEDGEMENTS

We would like to thank members of the Chapman lab and K. Bjornson for insightful comments and valuable discussion related to the manuscript. This study was supported by grants from the NIH (MH061876 and NS097362 to E.R.C.). C.S.E. was supported in part by a PhRMA Foundation predoctoral fellowship and by the UW-Madison Molecular and Cellular Pharmacology Training Grant (T32 GM008688). E.R.C. is an Investigator of the Howard Hughes Medical Institute.

This article is subject to HHMI’s Open Access to Publications policy. HHMI lab heads have previously granted a nonexclusive CC BY 4.0 license to the public and a sublicensable license to HHMI in their research articles. Pursuant to those licenses, the author-accepted manuscript of this article can be made freely available under a CC BY 4.0 license immediately upon publication.

## AUTHOR CONTRIBUTIONS

M.J.S.: conceptualization, data curation, formal analysis, investigation, methodology, project administration, software, supervision, validation, visualization, writing – original draft, writing – review and editing. C.S.E.: investigation, formal analysis, visualization. K.S.S.: investigation, formal analysis, validation, visualization. Z.W.: investigation, formal analysis, validation. E.R.C.: conceptualization, funding acquisition, methodology, project administration, resources, supervision, visualization, writing – original draft, writing – review and editing.

## SUPPLEMENTARY MATERIALS

**Supplementary Table 1.** Thermodynamic properties of Ca^2+^ binding to the isolated C2A and C2B domains of SYT9 as determined using ITC. Representative traces of SYT9 C2-domain•Ca^2+^ titrations are shown in (Fig. 6c). Data presented as mean ± SEM, *n* = 4.

|  | K<sub>D</sub> (mM) | DH (cal/mol) | DS (cal/mol/K) | DG (kcal/mol) |
| --- | --- | --- | --- | --- |
| SYT9 C2A | K<sub>D1</sub> = 53.0 ± 8.6 | DH<sub>1</sub> = 285 ± 65 | DS<sub>1</sub> = 20.6 ± 0.49 | DG<sub>1</sub> = -5.85 ± 0.08 |
|  | K<sub>D2</sub> = 182 ± 39 | DH<sub>2</sub> = 230 ± 220 | DS<sub>2</sub> = 18.0 ± 1.1 | DG<sub>2</sub> = -5.14 ± 0.12 |
|  | K<sub>D3</sub> = 337 ± 80 | DH<sub>3</sub> = 701 ± 250 | DS<sub>3</sub> = 18.5 ± 0.50 | DG<sub>3</sub> = -4.79 ± 0.15 |
| SYT9 C2B | K<sub>D1</sub> = 42.9 ± 5.5 | DH<sub>1</sub> = -403 ± 20 | DS<sub>1</sub> = 18.7 ± 0.30 | DG<sub>1</sub> = -5.97 ± 0.07 |
|  | K<sub>D2</sub> = 500 ± 95 | DH<sub>2</sub> = -878 ± 78 | DS<sub>2</sub> = 12.3 ± 0.58 | DG<sub>2</sub> = -4.53 ± 0.10 |

**Supplementary Figure 1.**
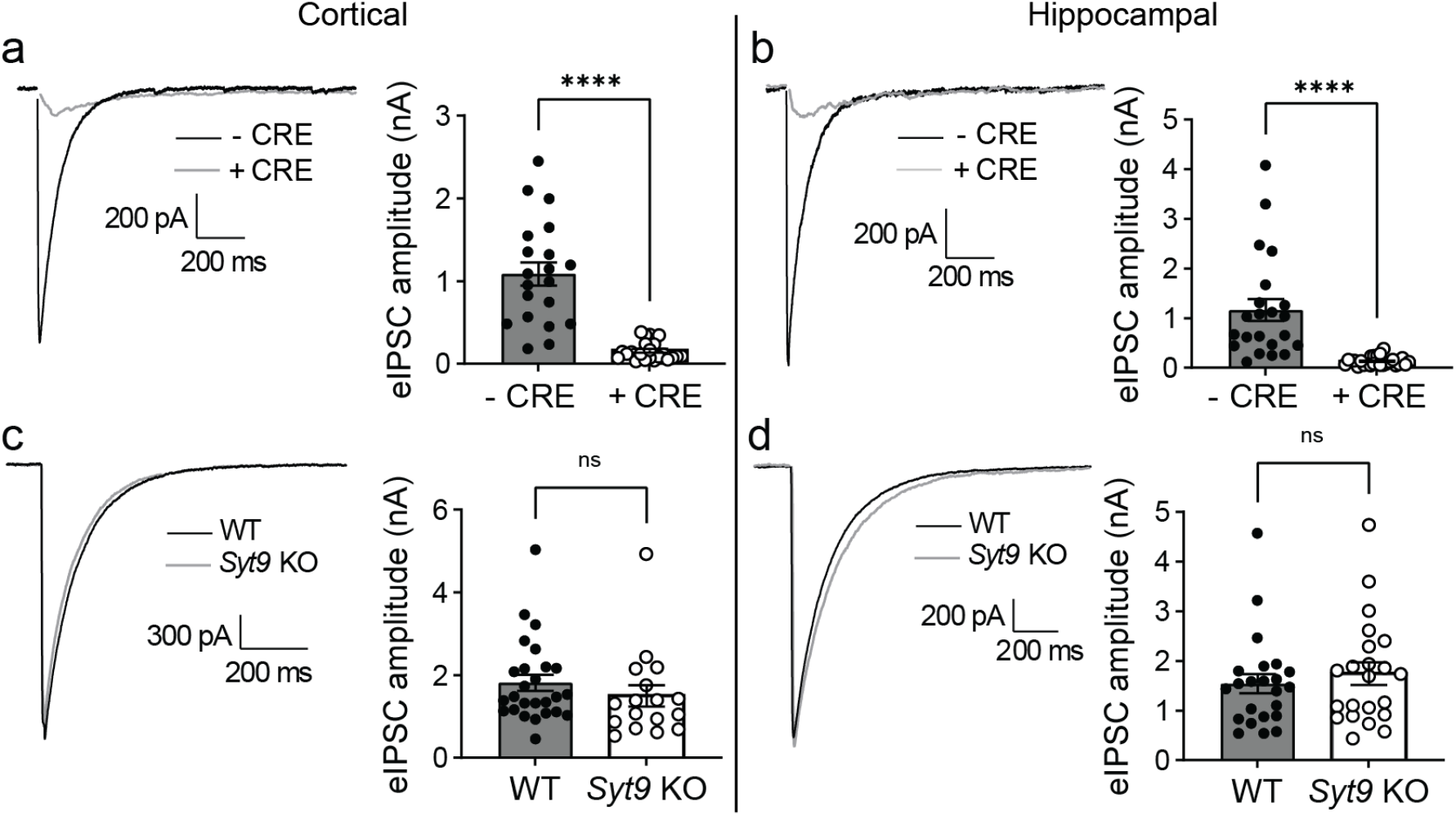
SYT1, but not SYT9, supports evoked neurotransmitter release in cortical and hippocampal neurons. **a**-**b**, Averaged eIPSC traces recorded from WT and *Syt1* cKO cortical (- CRE n = 26; + CRE n = 17) (**a**) and hippocampal (- CRE n = 23; + CRE n = 22) (**b**) neurons with quantification of peak eIPSC amplitude. **c**-**d**, Same as (**a-b**), but using *Syt9* KO neurons; cortical neurons (WT n = 20; KO n = 23); hippocampal neurons (WT n = 22; KO n = 26). eIPSCs were unaffected in *Syt9 KO* cortical (p = 0.3231; unpaired t-test) and hippocampal neurons (p = 0.4966; unpaired t-test). In contrast, eIPSCs were disrupted in *Syt1* cKO + CRE cortical (p < 0.0001; unpaired t-test) and hippocampal neurons (p < 0.0001; unpaired t-test).

**Supplementary Figure 2.**
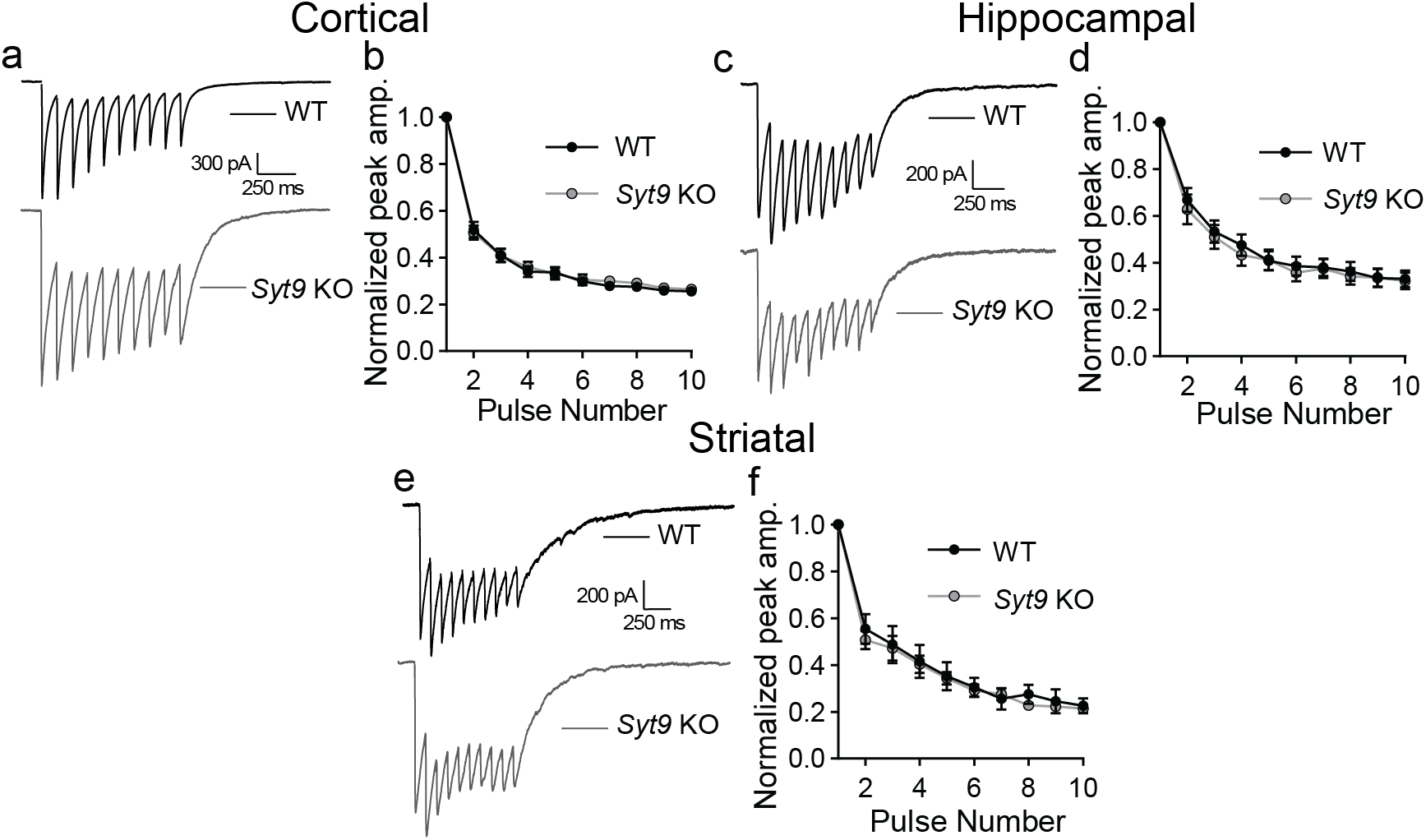
Synaptic depression was unchanged in *Syt9* KO cortical, hippocampal, and striatal neurons. **a**, Averaged 10 Hz eIPSC traces from WT (n = 21) and *Syt9* KO (n = 21) cortical neurons. **b**, Quantification of peak amplitudes from each pulse normalized to the first pulse, showing no difference in short-term plasticity. **c**-**d**, Same as **a**-**b**, but with hippocampal neurons (WT n = 17; KO n = 17). **e**-**f**, Same as **a**-**b**, but with striatal neurons (WT n = 35; KO n = 38). Error bars indicate SEM.

**Supplementary Figure 3.**
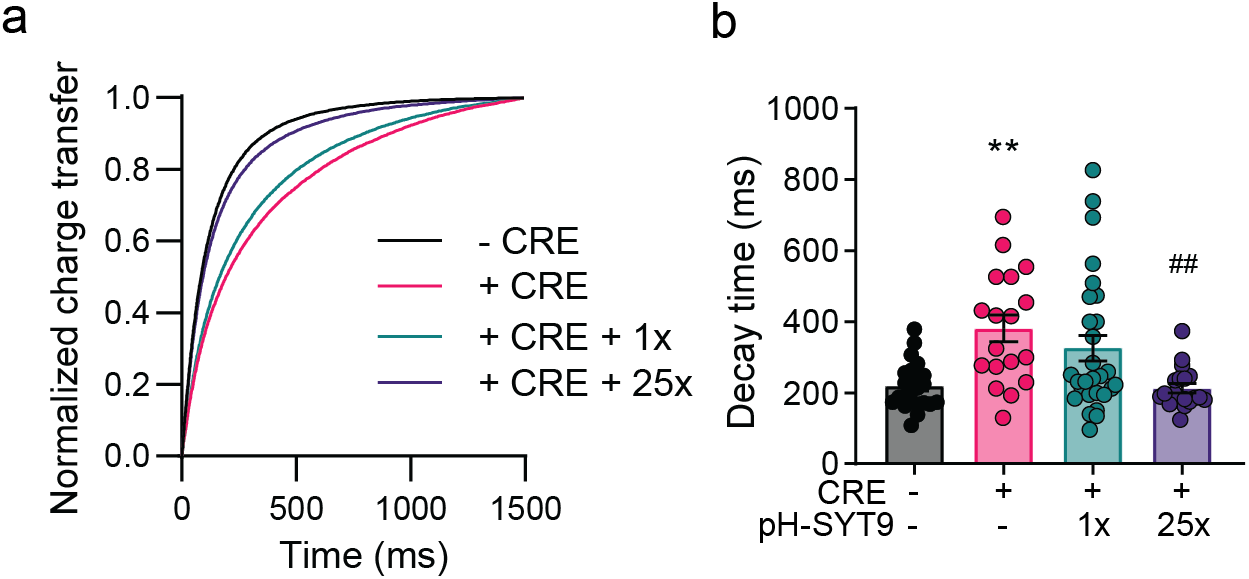
eIPSC charge transfer and decay kinetics recorded from *Syt1* cKO cortical neurons expressing pH-SYT9. **a**, Plot of normalized cumulative charge transfer of - CRE, + CRE, + CRE + 1x pH-SYT9, and + CRE + 25x pH-SYT9 conditions. **b**, Plot of mean decay time (90%-10%) for each condition; 25x fully rescues the decay time, while 1x fails to rescue. Further decay analysis was not performed due to receptor-dominated decay kinetics of IPSCs in cultured neurons. In this figure, asterisks (*) indicate statistical comparisons (one-way ANOVA with Šídák’s multiple comparisons test) to *Syt1* cKO neurons without CRE; hash symbols (#) indicate statistical comparisons to *Syt1* cKO neurons + CRE, where ##p < 0.01.

**Supplementary Figure 4.**
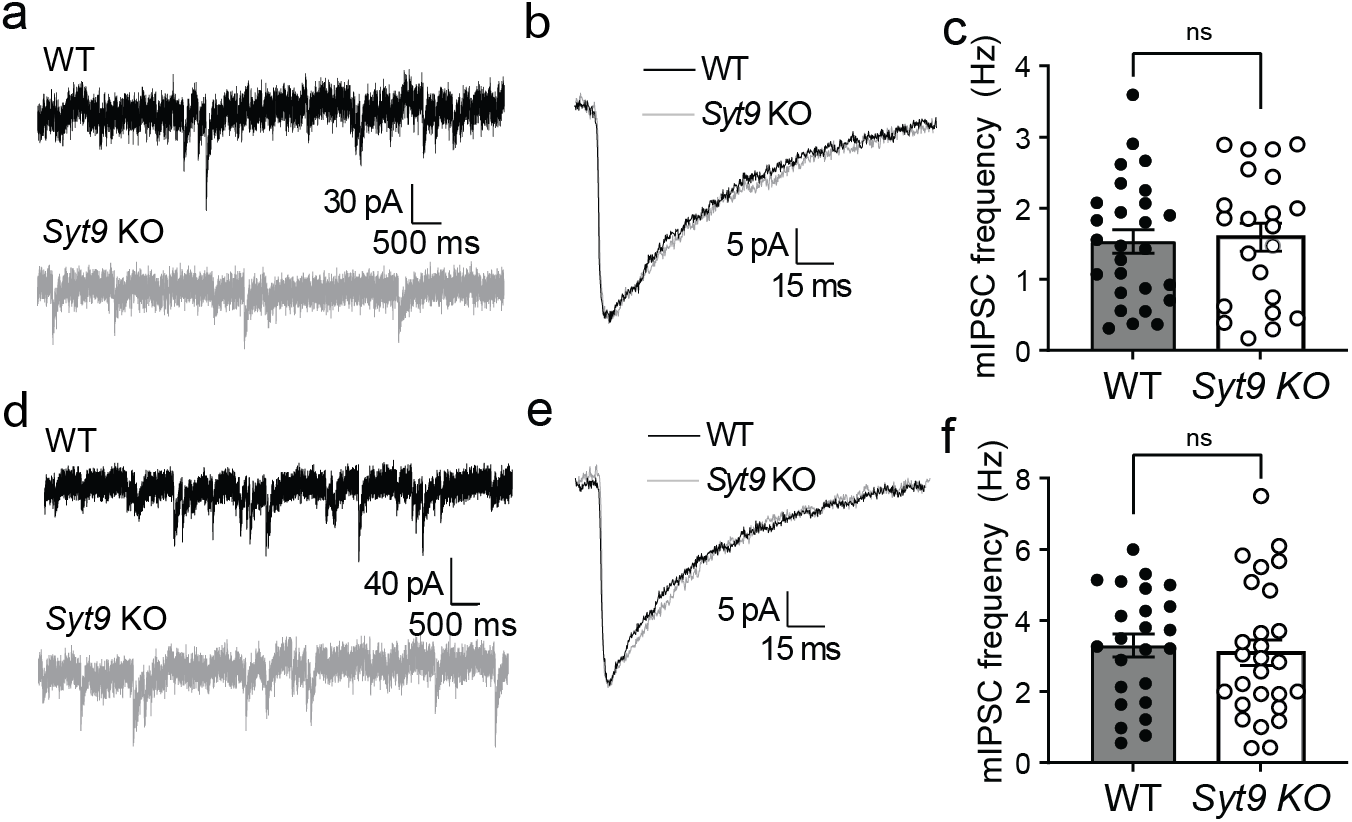
mIPSC frequency is unaltered in cultured *Syt9* KO cortical and hippocampal neurons. **A**, Representative mIPSC traces recorded from WT (n = 27) and *Syt9* KO (n = 22) cortical neurons. **B**, Overlay of averaged mIPSCs, revealing no apparent change in kinetics or amplitude. **C**, mIPSC frequency was unaltered in *Syt9* KO cortical neurons (p = 0.8181; unpaired t-test) **d**-**f**, Same as **a**-**c**, but using hippocampal neurons (WT n = 24; KO n = 27) (p = 0.6836; unpaired t-test).

**Supplementary Figure 5.**
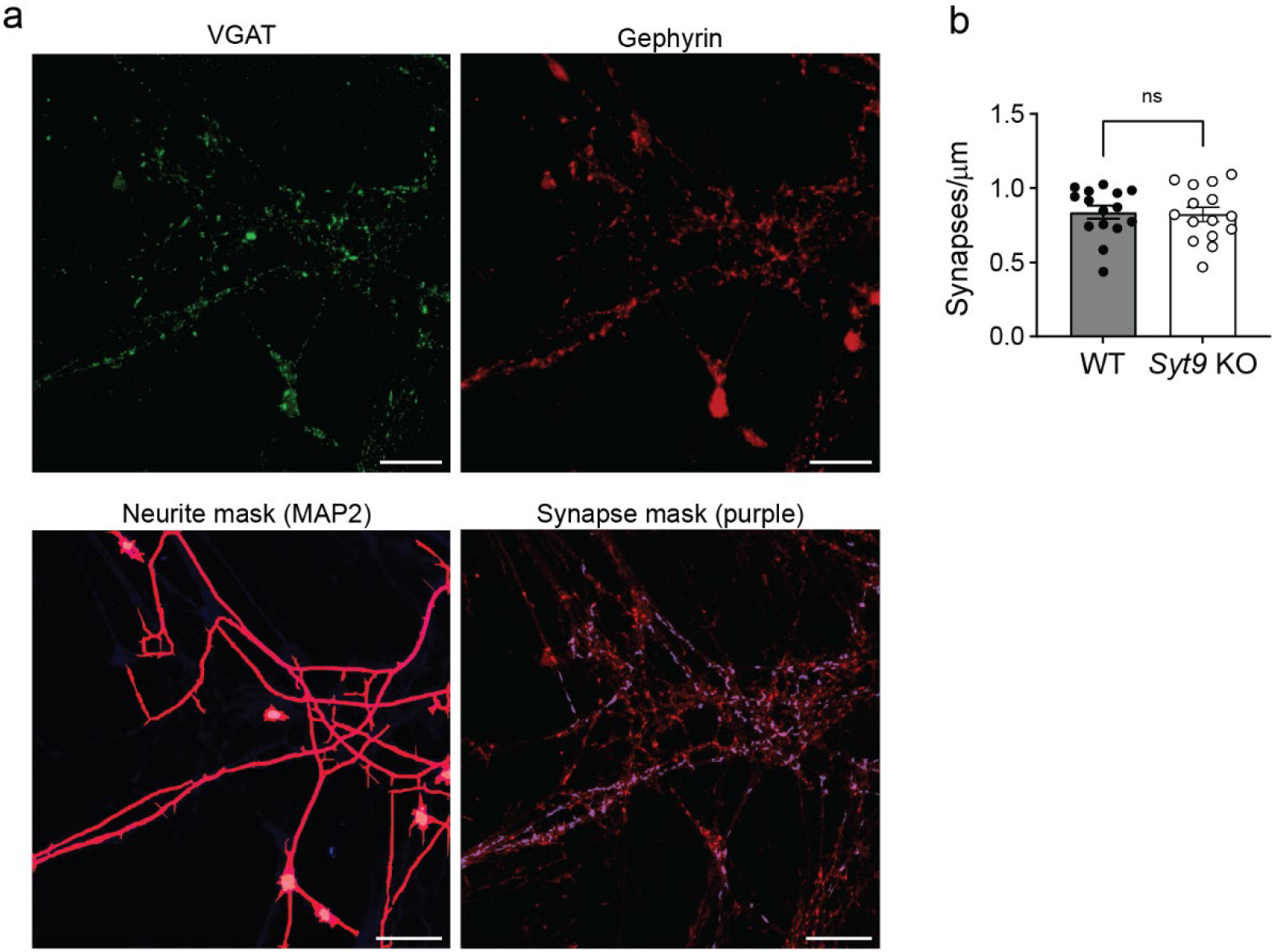
Unchanged synaptic density in *Syt9* KO striatal neurons. **a**, Representative images of cultured striatal neurons stained with α-Gephyrin (red) and α-VGAT (green) antibodies with a neurite mask generated from MAP2 staining (magenta). Gephyrin and VMAT overlap within the neurite mask were counted (purple puncta within synapse mask). Scale bars represent 20 μm. **b**, Number of synapses per μm is unaltered in cultured *Syt9* striatal neurons (n = 15) compared to WT neurons (n = 15) (p = 0.8212; unpaired t-test).

**Supplementary Figure 6.**
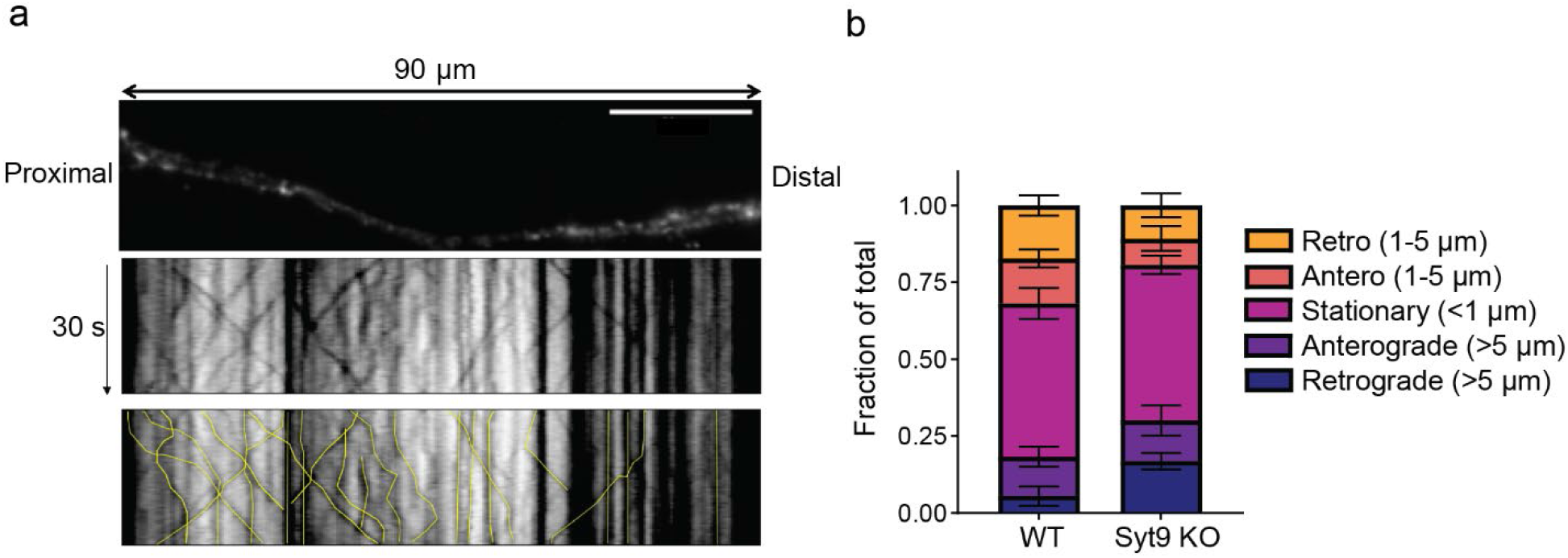
The trafficking of SP-pH-bearing DCVs is unaltered in *Syt9* KO striatal neurons. **a**, *Upper panel*: Representative image of a 90 µm region of a striatal neuron transfected with SP-pH; image was captured after perfusion with NH_4_Cl. *Lower panels*: Color inverted kymographs of this region, over a 30 s time period; each track in the kymograph was traced and quantified in the bottom panel (yellow lines). **b**, Quantification of the displacement and direction of SP-pH vesicles. No significant differences in WT (n = 98 vesicles, from 5 cells) versus *Syt9* KO (n = 101 vesicles, from 5 cells) striatal neurons were detected.

**Supplementary Figure 7.**
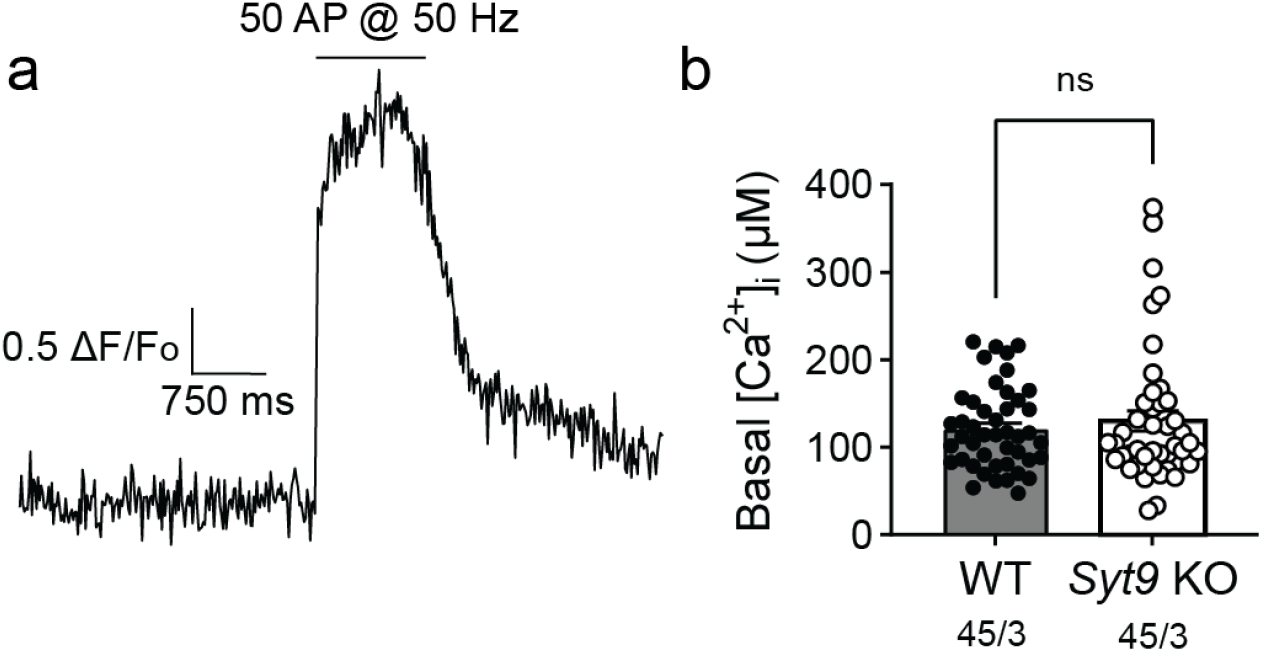
Basal presynaptic [Ca^2+^]_i_ is unchanged in *Syt9* KO striatal neurons. **a**, In each trial, the baseline signal was recorded for 3 s, followed by high-frequency train stimulation to obtain the maximal fluorescence of the indicator, to calculate the basal [Ca^2+^]_i_ (see Methods). **b**, Basal presynaptic [Ca^2+^]_i_ in 1.5 mM [Ca^2+^]_e_ did not differ between WT (n = 45) and *Syt9* KO (n = 44) striatal neurons (p = 0.8735; Mann-Whitney test).

**Supplementary Figure 8.**
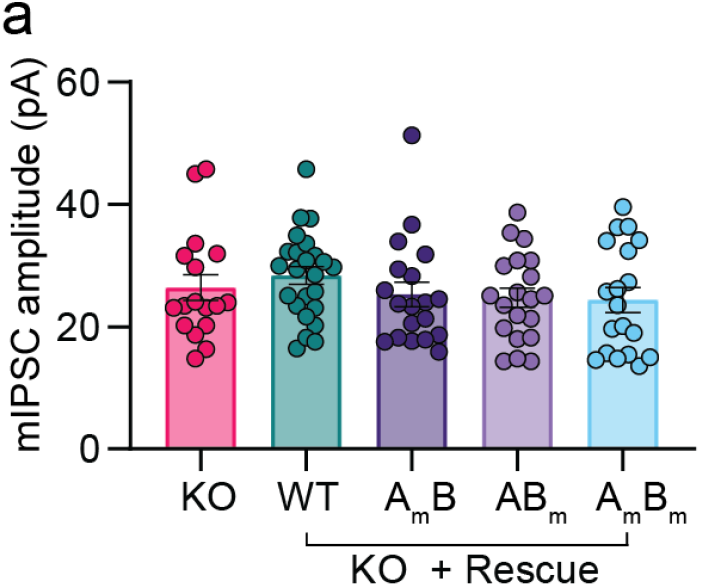
mIPSC amplitude is unaltered in SYT9 CLM rescue. **a**, mIPSC amplitudes measured from (Fig. 7). No significant differences in average amplitudes between conditions were measured (p = 0.4169; Kruskal Wallis test).

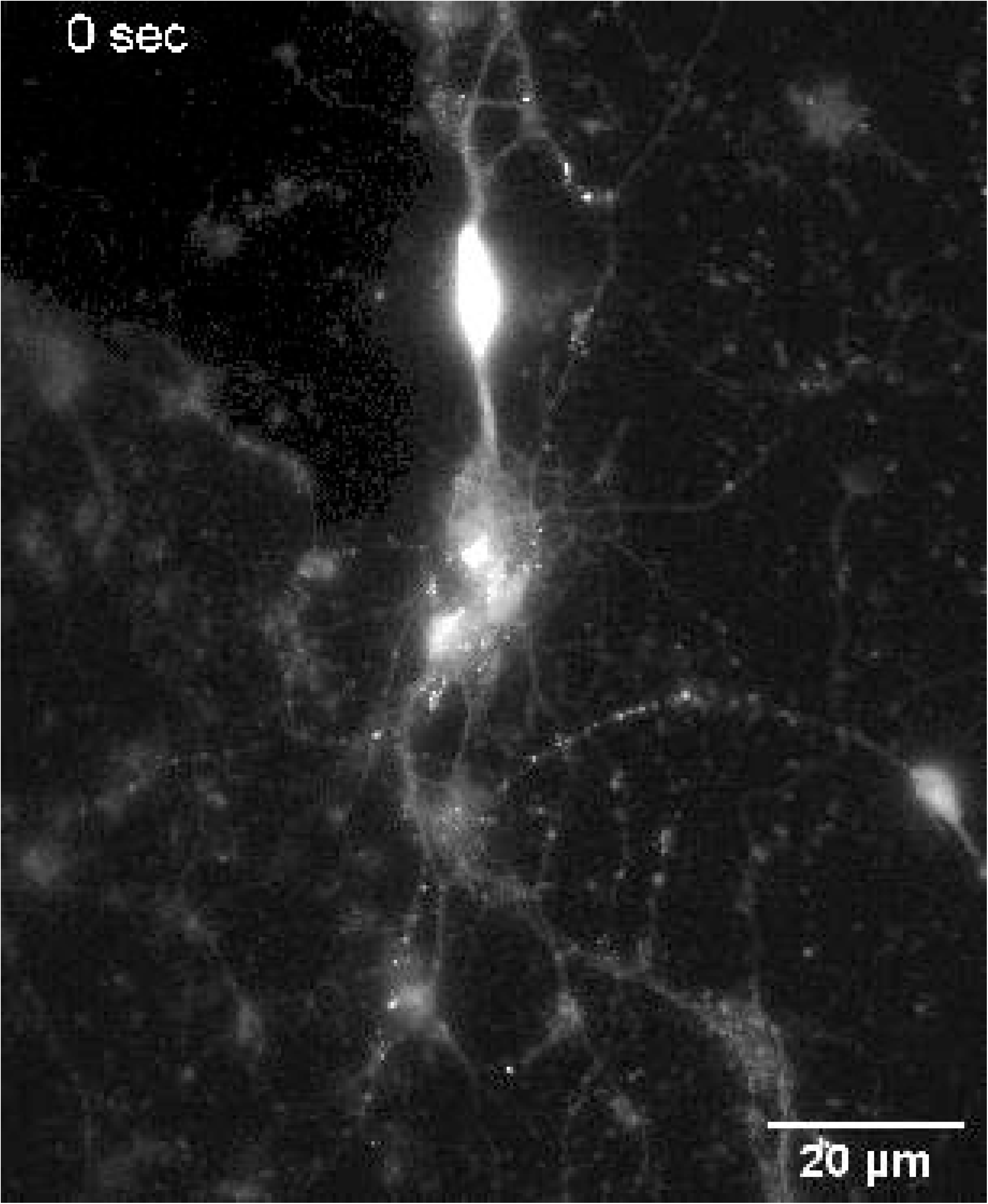

