## Supplementary material for "Synaptotagmin 9 modulates spontaneous neurotransmitter release in striatal neurons by regulating substance P secretion": Video 1 Legend

**SYT9 regulates the secretion of SP-pH from striatal neurons**

Representative timelapse of SP-pH release events in a WT striatal neuron during electrical stimulation (16 x [50 AP at 50 Hz]). Stimulation starts at t = 10 s and ends at t = 34 s; $NH_4$ perfusion started at t = 80 s.
