## Supplementary material for "Synaptotagmin 9 modulates spontaneous neurotransmitter release in striatal neurons by regulating substance P secretion": Source data and statistics

**Figure 1b**

| | Ctx | Hip | Str |
|---|---|---|---|
| Number of values | 6 | 6 | 6 |
| Mean (percentage) | 0.100 | 0.121 | 0.0628 |
| Median | 0.0838 | 0.107 | 0.0651 |
| Std. Deviation | 0.0373 | 0.0464 | 0.0168 |
| Std. Error of Mean | 0.0152 | 0.0190 | 0.00687 |

**Figure 1d**

| | WT | *Syt1* cKO |
|---|---|---|
| Number of values | 21 | 22 |
| Mean (nA) | 0.854 | 0.126 |
| Median | 0.641 | 0.111 |
| Std. Deviation | 0.665 | 0.0732 |
| Std. Error of Mean | 0.145 | 0.0156 |

| | |
|---|---|
| Mann Whitney test | |
| P value | <0.0001 |
| P value summary | **** |
| Significantly different (P < 0.05)? | Yes |
| One- or two-tailed P value? | Two-tailed |
| Sum of ranks in column A,B | 684 , 262 |
| Mann-Whitney U | 9 |

**Figure 1f**

| | Ctx | Hip | Str |
|---|---|---|---|
| Number of values | 4 | 4 | 4 |
| Mean (percentage) | 0.00373 | 0.00442 | 0.00310 |
| Median | 0.00374 | 0.00452 | 0.00290 |
| Std. Deviation | 0.000143 | 0.000396 | 0.000904 |
| Std. Error of Mean | 7.14e-005 | 0.000198 | 0.000452 |

**Figure 1h**

| | WT | *Syt9* KO |
|---|---|---|
| Number of values | 28 | 27 |
| Mean (nA) | 1.03 | 0.959 |
| Median | 0.911 | 0.899 |
| Std. Deviation | 0.676 | 0.626 |
| Std. Error of Mean | 0.128 | 0.121 |

| Unpaired t test | |
|---|---|
| P value | 0.6979 |
| P value summary | ns |
| Significantly different (P < 0.05)? | No |
| One- or two-tailed P value? | Two-tailed |
| t, df | t=0.3903, df=53 |

**Figure 2c**

| | - Cre | + Cre | + Cre + 1x pH-Syt9 | + Cre + 25x pH-Syt9 |
|---|---|---|---|---|
| Number of values | 21 | 19 | 25 | 23 |
| Mean (pA) | 1773 | 122 | 257 | 1035 |
| Median | 1747 | 117 | 161 | 849 |
| Std. Deviation | 1129 | 49.7 | 187 | 570 |
| Std. Error of Mean | 246 | 11.4 | 37.5 | 119 |

| **Kruskal-Wallis test** | |
|---|---|
| P value | <0.0001 |
| P value summary | **** |
| Do the medians vary signif. (P < 0.05)? | Yes |
| Number of groups | 4 |
| Kruskal-Wallis statistic | 64.41 |

| **Dunn's multiple comparisons test** | Mean rank diff. | Significant? | Summary | Adjusted P Value |
|---|---|---|---|---|
| - Cre vs. + Cre | 54.63 | Yes | **** | <0.0001 |
| - Cre vs. + Cre + 1x pH-Syt9 | 42.37 | Yes | **** | <0.0001 |
| - Cre vs. + Cre + 25x pH-Syt9 | 10.40 | No | ns | 0.8880 |
| + Cre vs. + Cre + 1x pH-Syt9 | -12.26 | No | ns | 0.5744 |
| + Cre vs. + Cre + 25x pH-Syt9 | -44.23 | Yes | **** | <0.0001 |

**Figure 2d**

| | - Cre | + Cre | + Cre + 1x pH-Syt9 | + Cre + 25x pH-Syt9 |
|---|---|---|---|---|
| Number of values | 22 | 19 | 27 | 24 |
| Mean (ms) | 6.90 | 37.9 | 22.1 | 12.5 |
| Median | 6.30 | 36.6 | 21.0 | 13.8 |
| Std. Deviation | 1.54 | 19.3 | 13.5 | 5.84 |
| Std. Error of Mean | 0.329 | 4.44 | 2.61 | 1.19 |

| **Kruskal-Wallis test** | |
|---|---|
| P value | <0.0001 |
| P value summary | **** |
| Do the medians vary signif. (P < 0.05)? | Yes |

| | |
|---|---|
| Number of groups | 4 |
| Kruskal-Wallis statistic | 41.41 |

| **Dunn's multiple comparisons test** | Mean rank diff. | Significant? | Summary | Adjusted P Value |
|---|---|---|---|---|
| - Cre vs. + Cre | -50.97 | Yes | **** | <0.0001 |
| - Cre vs. + Cre + 1x pH-Syt9 | -32.99 | Yes | **** | <0.0001 |
| - Cre vs. + Cre + 25x pH-Syt9 | -17.84 | No | ns | 0.1178 |
| + Cre vs. + Cre + 1x pH-Syt9 | 17.98 | No | ns | 0.1227 |
| + Cre vs. + Cre + 25x pH-Syt9 | 33.13 | Yes | *** | 0.0003 |

**Figure 2f**

| | - Cre | + Cre | + Cre + 1x pH-Syt9 | +Cre + 25x pH-Syt9 |
|---|---|---|---|---|
| Number of values | 12 | 10 | 16 | 15 |
| Mean (Hz) | 0.976 | 2.89 | 3.30 | 1.63 |
| Median | 1.04 | 2.88 | 3.46 | 1.79 |
| Std. Deviation | 0.397 | 1.24 | 0.999 | 0.689 |
| Std. Error of Mean | 0.115 | 0.392 | 0.250 | 0.178 |

| ANOVA summary | |
|---|---|
| F | 20.65 |
| P value | <0.0001 |
| P value summary | **** |
| Significant diff. among means (P < 0.05)? | Yes |
| R squared | 0.5584 |

| Šídák's multiple comparisons test | Mean Diff. | 95.00% CI of diff. | Below threshold? | Summary | Adjusted P Value |
|---|---|---|---|---|---|
| - Cre vs. + Cre | -1.916 | -2.913 to -0.9195 | Yes | **** | <0.0001 |
| - Cre vs. + Cre + 1x pH-Syt9 | -2.323 | -3.212 to -1.434 | Yes | **** | <0.0001 |
| - Cre vs. +Cre + 25x pH-Syt9 | -0.6498 | -1.551 to 0.2516 | No | ns | 0.2657 |
| + Cre vs. + Cre + 1x pH-Syt9 | -0.4068 | -1.345 to 0.5315 | No | ns | 0.7663 |
| + Cre vs. +Cre + 25x pH-Syt9 | 1.266 | 0.3160 to 2.216 | Yes | ** | 0.0042 |

**Figure 2h**

| | - Cre | + Cre | + Cre + 1x pH-Syt9 | +Cre + 25x pH-Syt9 |
|---|---|---|---|---|
| Number of values | 11 | 10 | 16 | 15 |
| Mean (pA) | 42.0 | 41.8 | 42.9 | 42.1 |
| Median | 42.1 | 42.9 | 44.6 | 45.0 |
| Std. Deviation | 5.93 | 6.61 | 7.21 | 8.92 |
| Std. Error of Mean | 1.79 | 2.09 | 1.80 | 2.30 |

| ANOVA summary | |
|---|---|
| F | 0.05727 |
| P value | 0.9818 |
| P value summary | ns |
| Significant diff. among means (P < 0.05)? | No |
| R squared | 0.003567 |

**Figure 2i**

| | 1x (internal) | 25x (internal) | Endogenous SYT1 | Endogenous SYT9 |
|---|---|---|---|---|
| Number of values | 25 | 21 | 18 | 18 |
| Mean (PCC) | 0.618 | 0.747 | 0.752 | 0.597 |
| Median | 0.610 | 0.740 | 0.770 | 0.605 |
| Std. Deviation | 0.0561 | 0.0222 | 0.0260 | 0.0390 |
| Std. Error of Mean | 0.0112 | 0.00484 | 0.00613 | 0.00918 |

| ANOVA summary | |
|---|---|
| F | 86.69 |
| P value | <0.0001 |
| P value summary | **** |
| Significant diff. among means (P < 0.05)? | Yes |
| R squared | 0.7693 |

| Tukey's multiple comparisons test | Mean Diff. | 95.00% CI of diff. | Below threshold? | Summary | Adjusted P Value |
|---|---|---|---|---|---|
| 1x (int.) vs. 25x (int.) | -0.1295 | -0.1604 to -0.09871 | Yes | **** | <0.0001 |
| 1x (int.) vs. Endo SYT1 | -0.1346 | -0.1668 to -0.1024 | Yes | **** | <0.0001 |
| 1x (int.) vs. Endo SYT9 | 0.02093 | -0.01127 to 0.05313 | No | ns | 0.3272 |
| 25x (int.) vs. Endo SYT1 | -0.005079 | -0.03854 to 0.02838 | No | ns | 0.9784 |
| 25x (int.) vs. Endo SYT9 | 0.1505 | 0.1170 to 0.1839 | Yes | **** | <0.0001 |
| Endo SYT1 vs. Endo SYT9 | 0.1556 | 0.1208 to 0.1903 | Yes | **** | <0.0001 |

**Figure 3c**

| | WT | *Syt9* KO | *Syt9* KO + WT rescue |
|---|---|---|---|
| Number of values | 33 | 29 | 24 |
| Mean (Hz) | 1.01 | 0.432 | 1.15 |
| Median | 0.973 | 0.357 | 1.01 |
| Std. Deviation | 0.571 | 0.296 | 0.505 |

| | | | |
|---|---|---|---|
| Std. Error of Mean | 0.0994 | 0.0550 | 0.103 |

| ANOVA summary | |
|---|---|
| F | 17.73 |
| P value | <0.0001 |
| P value summary | **** |
| Significant diff. among means (P < 0.05)? | Yes |
| R squared | 0.2994 |

| Tukey's multiple comparisons test | Mean Diff. | 95.00% CI of diff. | Below threshold? | Summary | Adjusted P Value |
|---|---|---|---|---|---|
| WT vs. syt9 KO | 0.5775 | 0.2886 to 0.8663 | Yes | **** | <0.0001 |
| WT vs. syt9 KO + WT rescue | -0.1373 | -0.4417 to 0.1671 | No | ns | 0.5314 |
| syt9 KO vs. syt9 KO + WT rescue | -0.7147 | -1.028 to -0.4016 | Yes | **** | <0.0001 |

**Figure 4b**

| | Syt9: SubP-pH | SubP-pH: Syt9 |
|---|---|---|
| Number of values | 23 | 21 |
| Mean (MOC) | 0.655 | 0.808 |
| Median | 0.641 | 0.801 |
| Std. Deviation | 0.123 | 0.0799 |
| Std. Error of Mean | 0.0257 | 0.0174 |

**Figure 4d**

| | ChgB: SubP-pH | SubP-pH: ChgB |
|---|---|---|
| Number of values | 10 | 10 |
| Mean (MOC) | 0.855 | 0.881 |
| Median | 0.849 | 0.875 |
| Std. Deviation | 0.0374 | 0.0239 |
| Std. Error of Mean | 0.0118 | 0.00755 |

**Figure 4g**

| | WT | Syt9 KO |
|---|---|---|
| Number of values | 21 | 26 |
| Mean (fraction) | 0.103 | 0.0398 |

| | | |
|---|---|---|
| Median | 0.0776 | 0.0204 |
| Std. Deviation | 0.0867 | 0.0472 |
| Std. Error of Mean | 0.0189 | 0.00926 |

| Mann Whitney test | |
|---|---|
| P value | 0.0049 |
| Exact or approximate P value? | Exact |
| P value summary | ** |
| Significantly different (P < 0.05)? | Yes |
| One- or two-tailed P value? | Two-tailed |
| Sum of ranks in column A,B | 633.5 , 494.5 |
| Mann-Whitney U | 143.5 |

**Figure 4h**

| | WT | Syt9 KO |
|---|---|---|
| Number of values | 21 | 26 |
| Mean (# vesicles) | 656.9 | 731.3 |
| Median | 489.0 | 565.5 |
| Std. Deviation | 548.7 | 578.0 |
| Std. Error of Mean | 119.7 | 113.4 |

| Mann Whitney test | |
|---|---|
| P value | 0.6034 |
| Exact or approximate P value? | Exact |
| P value summary | ns |
| Significantly different (P < 0.05)? | No |
| One- or two-tailed P value? | Two-tailed |
| Sum of ranks in column A,B | 479 , 649 |
| Mann-Whitney U | 248 |

**Figure 4j**

| | WT | Syt9 KO |
|---|---|---|
| Number of values | 1749 | 1111 |
| Mean (s) | 5.471 | 12.25 |
| Median | 1.000 | 1.800 |
| Std. Deviation | 12.06 | 20.45 |
| Std. Error of Mean | 0.2883 | 0.6135 |

**Figure 5b**

| | WT | KO | KO + SubP | KO + GR7 | WT + SR1 | WT + SubP | KO + L-Enk | KO + Dyn A | KO + CCK8 | KO + SR1 |
|---|---|---|---|---|---|---|---|---|---|---|
| Number of values | 27 | 39 | 35 | 26 | 34 | 28 | 23 | 23 | 20 | 27 |
| Mean (Hz) | 0.931 | 0.430 | 1.40 | 0.967 | 0.578 | 1.07 | 0.374 | 0.466 | 0.450 | 0.564 |
| Median | 0.889 | 0.350 | 1.00 | 0.823 | 0.514 | 0.995 | 0.307 | 0.417 | 0.417 | 0.517 |
| Std. Deviation | 0.369 | 0.240 | 0.859 | 0.611 | 0.284 | 0.393 | 0.231 | 0.346 | 0.288 | 0.413 |
| Std. Error of Mean | 0.0709 | 0.0385 | 0.145 | 0.120 | 0.0487 | 0.0743 | 0.0482 | 0.0722 | 0.0645 | 0.0795 |

| Kruskal-Wallis test | |
|---|---|
| P value | <0.0001 |
| Exact or approximate P value? | Approximate |
| P value summary | **** |
| Do the medians vary signif. (P < 0.05)? | Yes |
| Number of groups | 10 |
| Kruskal-Wallis statistic | 111.0 |

| Dunn's multiple comparisons test | Mean rank diff. | Significant? | Summary | Adjusted P Value |
|---|---|---|---|---|
| WT vs. KO | 87.29 | Yes | **** | <0.0001 |
| WT vs. KO + SubP | -23.55 | No | ns | >0.9999 |
| WT vs. KO + GR73632 | 5.074 | No | ns | >0.9999 |
| WT vs. WT + SR140333 | 56.38 | Yes | * | 0.0322 |
| WT vs. WT + SubP | -17.65 | No | ns | >0.9999 |
| WT vs. KO + L-Enk | 98.62 | Yes | **** | <0.0001 |
| WT vs. KO + Dyn A | 79.36 | Yes | ** | 0.0013 |
| WT vs. KO + CCK8 | 83.46 | Yes | ** | 0.0011 |
| WT vs. KO + SR140333 | 74.57 | Yes | * | 0.0133 |
| KO vs. KO + SubP | -110.8 | Yes | **** | <0.0001 |
| KO vs. KO + GR73632 | -82.22 | Yes | *** | 0.0005 |
| KO vs. WT + SR140333 | -30.91 | No | ns | >0.9999 |
| KO vs. WT + SubP | -104.9 | Yes | **** | <0.0001 |
| KO vs. KO + L-Enk | 11.33 | No | ns | >0.9999 |
| KO vs. KO + Dyn A | -7.928 | No | ns | >0.9999 |
| KO vs. KO + CCK8 | -3.835 | No | ns | >0.9999 |
| KO vs. KO + SR140333 | -25.86 | No | ns | >0.9999 |

**Figure 5c**

| | WT | KO | KO + SubP | KO + GR7 | WT + SR1 | WT + SubP | KO + L-Enk | KO + Dyn A | KO + CCK8 | KO + SR1 |
|---|---|---|---|---|---|---|---|---|---|---|
| Number of values | 35 | 39 | 35 | 26 | 28 | 28 | 22 | 23 | 20 | 27 |
| Mean (pA) | 22.7 | 22.8 | 21.0 | 24.1 | 24.5 | 21.5 | 24.3 | 22.4 | 20.2 | 23.3 |
| Median | 21.6 | 20.6 | 17.8 | 23.2 | 24.0 | 18.9 | 24.6 | 24.1 | 18.3 | 23.4 |
| Std. Deviation | 8.03 | 7.38 | 10.0 | 6.36 | 5.73 | 9.28 | 5.12 | 6.22 | 6.41 | 5.18 |

| | | | | | | | | | | |
|---|---|---|---|---|---|---|---|---|---|---|
| Std. Error of Mean | 1.36 | 1.18 | 1.69 | 1.50 | 1.12 | 1.75 | 1.09 | 1.30 | 1.43 | 0.996 |

| Kruskal-Wallis test | |
|---|---|
| P value | 0.0562 |
| Exact or approximate P value? | Approximate |
| P value summary | ns |
| Do the medians vary signif. (P < 0.05)? | No |
| Number of groups | 10 |
| Kruskal-Wallis statistic | 16.55 |

**Figure 7a**

| | KO | KO+WT | KO+C2AmB | KO+C2ABm | KO+C2AmBm |
|---|---|---|---|---|---|
| Number of values | 24 | 23 | 22 | 24 | 19 |
| Mean (fraction) | 0.0394 | 0.0889 | 0.0382 | 0.110 | 0.0356 |
| Median | 0.0343 | 0.0756 | 0.0131 | 0.0893 | 0.0213 |
| Std. Deviation | 0.0318 | 0.0635 | 0.0525 | 0.0917 | 0.0310 |
| Std. Error of Mean | 0.00650 | 0.0132 | 0.0112 | 0.0187 | 0.00712 |

| Kruskal-Wallis test | |
|---|---|
| P value | <0.0001 |
| Exact or approximate P value? | Approximate |
| P value summary | **** |
| Do the medians vary signif. (P < 0.05)? | Yes |
| Number of groups | 5 |
| Kruskal-Wallis statistic | 27.65 |

| Dunn's multiple comparisons test | Mean rank diff. | Significant? | Summary | Adjusted P Value |
|---|---|---|---|---|
| KO vs. KO+WT | -27.32 | Yes | * | 0.0158 |
| KO vs. KO+C2AmB | 8.998 | No | ns | >0.9999 |
| KO vs. KO+C2ABm | -29.38 | Yes | ** | 0.0069 |
| KO vs. KO+C2AmBm | 2.402 | No | ns | >0.9999 |

**Figure 7b**

| | KO | KO+WT | KO+C2AmB | KO+C2ABm | KO+C2AmBm |
|---|---|---|---|---|---|
| Number of values | 24 | 23 | 22 | 24 | 19 |
| Mean (# vesicles) | 796 | 796 | 763 | 650 | 789 |
| Median | 704 | 684 | 612 | 540 | 698 |
| Std. Deviation | 491 | 412 | 552 | 415 | 420 |
| Std. Error of Mean | 100 | 86.0 | 118 | 84.6 | 99.0 |

| Kruskal-Wallis test |
|---|

| | |
|---|---|
| P value | 0.6272 |
| Exact or approximate P value? | Approximate |
| P value summary | ns |
| Do the medians vary signif. (P < 0.05)? | No |
| Number of groups | 5 |
| Kruskal-Wallis statistic | 2.598 |

**Figure 7d**

| | KO | KO+WT | KO+C2AmB | KO+C2ABm | KO+C2AmBm |
|---|---|---|---|---|---|
| Number of values | 17 | 24 | 19 | 20 | 19 |
| Mean (Hz) | 0.493 | 1.15 | 0.487 | 1.17 | 0.531 |
| Median | 0.500 | 1.01 | 0.523 | 1.14 | 0.479 |
| Std. Deviation | 0.179 | 0.505 | 0.272 | 0.383 | 0.206 |
| Std. Error of Mean | 0.0434 | 0.103 | 0.0623 | 0.0856 | 0.0473 |

| Kruskal-Wallis test | |
|---|---|
| P value | <0.0001 |
| Exact or approximate P value? | Approximate |
| P value summary | **** |
| Do the medians vary signif. (P < 0.05)? | Yes |
| Number of groups | 5 |
| Kruskal-Wallis statistic | 57.16 |

| Dunn's multiple comparisons test | Mean rank diff. | Significant? | Summary | Adjusted P Value |
|---|---|---|---|---|
| KO vs. KO+WT | -43.15 | Yes | **** | <0.0001 |
| KO vs. KO+C2AmB | 0.1440 | No | ns | >0.9999 |
| KO vs. KO+C2ABm | -46.61 | Yes | **** | <0.0001 |
| KO vs. KO+C2AmBm | -2.777 | No | ns | >0.9999 |

**Supplementary Figure 1a**

| | WT | *Syt9* KO |
|---|---|---|
| Number of values | 20 | 23 |
| Mean | 1.09 | 0.158 |
| Median | 1.04 | 0.120 |
| Std. Deviation | 0.630 | 0.114 |
| Std. Error of Mean | 0.141 | 0.0237 |

| Unpaired t test | |
|---|---|
| P value | <0.0001 |
| P value summary | **** |
| Significantly different (P < 0.05)? | Yes |
| One- or two-tailed P value? | Two-tailed |

| | |
|---|---|
| t, df | t=6.957, df=41 |

**Supplementary Figure 1b**

| | WT | *Syt9* KO |
|---|---|---|
| Number of values | 22 | 26 |
| Mean | 1.16 | 0.131 |
| Median | 0.854 | 0.121 |
| Std. Deviation | 1.04 | 0.0869 |
| Std. Error of Mean | 0.221 | 0.0170 |

| | |
|---|---|
| Unpaired t test | |
| P value | <0.0001 |
| P value summary | **** |
| Significantly different (P < 0.05)? | Yes |
| One- or two-tailed P value? | Two-tailed |
| t, df | t=5.065, df=46 |

**Supplementary Figure 1c**

| | WT | *Syt9* KO |
|---|---|---|
| Number of values | 26 | 17 |
| Mean | 1.81 | 1.50 |
| Median | 1.48 | 1.31 |
| Std. Deviation | 0.985 | 1.05 |
| Std. Error of Mean | 0.193 | 0.255 |

| | |
|---|---|
| Unpaired t test | |
| P value | 0.3231 |
| P value summary | ns |
| Significantly different (P < 0.05)? | No |
| One- or two-tailed P value? | Two-tailed |
| t, df | t=1.000, df=41 |

**Supplementary Figure 1d**

| | WT | *Syt9* KO |
|---|---|---|
| Number of values | 23 | 22 |
| Mean | 1.54 | 1.74 |
| Median | 1.47 | 1.71 |
| Std. Deviation | 0.923 | 1.06 |
| Std. Error of Mean | 0.192 | 0.226 |

| |
|---|
| Unpaired t test |

| | |
|---|---|
| P value | 0.4966 |
| P value summary | ns |
| Significantly different (P < 0.05)? | No |
| One- or two-tailed P value? | Two-tailed |
| t, df | t=0.6856, df=43 |

**Supplementary Figure 3b**

| | - CRE | + CRE | + CRE + 1x pH-SYT9 | + CRE + 25x pH-SYT9 |
|---|---|---|---|---|
| Number of values | 22 | 19 | 28 | 21 |
| Mean | 218 | 361 | 325 | 256 |
| Median | 209 | 325 | 243 | 197 |
| Std. Deviation | 66.8 | 177 | 193 | 151 |
| Std. Error of Mean | 14.2 | 40.6 | 36.5 | 32.9 |

| **Kruskal-Wallis test** | |
|---|---|
| P value | 0.0078 |
| P value summary | ** |
| Do the medians vary signif. (P < 0.05)? | Yes |
| Number of groups | 4 |
| Kruskal-Wallis statistic | 11.87 |

| **Dunn's multiple comparisons test** | Mean rank diff. | Significant? | Summary | Adjusted P Value |
|---|---|---|---|---|
| - CRE vs. + CRE | -25.09 | Yes | * | 0.0108 |
| - CRE vs. + CRE + 1x pH-SYT9 | -15.42 | No | ns | 0.1915 |
| - CRE vs. + CRE + 25x pH-SYT9 | -3.693 | No | ns | >0.9999 |
| + CRE vs. + CRE + 25x pH-SYT9 | 21.39 | Yes | * | 0.0485 |
| + CRE vs. + CRE + 1x pH-SYT9 | 9.667 | No | ns | >0.9999 |

**Supplementary Figure 4c**

| | WT | *Syt9* KO |
|---|---|---|
| Number of values | 27 | 22 |
| Mean | 1.53 | 1.59 |
| Median | 1.47 | 1.80 |
| Std. Deviation | 0.865 | 0.933 |
| Std. Error of Mean | 0.166 | 0.199 |

| Unpaired t test | |
|---|---|
| P value | 0.8181 |
| P value summary | ns |
| Significantly different (P < 0.05)? | No |
| One- or two-tailed P value? | Two-tailed |

| | |
|---|---|
| t, df | t=0.2313, df=47 |

**Supplementary Figure 4f**

| | WT | *Syt9* KO |
|---|---|---|
| Number of values | 24 | 27 |
| Mean | 3.29 | 3.09 |
| Median | 3.38 | 2.84 |
| Std. Deviation | 1.59 | 1.89 |
| Std. Error of Mean | 0.325 | 0.363 |

| Unpaired t test | |
|---|---|
| P value | 0.6836 |
| P value summary | ns |
| Significantly different (P < 0.05)? | No |
| One- or two-tailed P value? | Two-tailed |
| t, df | t=0.4099, df=49 |

**Supplementary Figure 5b**

| | WT | *Syt9* KO |
|---|---|---|
| Number of values | 15 | 15 |
| Mean | 0.837 | 0.823 |
| Median | 0.872 | 0.812 |
| Std. Deviation | 0.168 | 0.184 |
| Std. Error of Mean | 0.0434 | 0.0476 |

| Unpaired t test | |
|---|---|
| P value | 0.8212 |
| P value summary | ns |
| Significantly different (P < 0.05)? | No |
| One- or two-tailed P value? | Two-tailed |
| t, df | t=0.2281, df=28 |

**Supplementary Figure 7b**

| | WT | Syt9 KO |
|---|---|---|
| Number of values | 45 | 44 |
| Mean | 121 | 130 |
| Median | 114 | 103 |
| Std. Deviation | 46.9 | 77.7 |

| | | |
|---|---|---|
| Std. Error of Mean | 6.99 | 11.7 |

| Mann Whitney test | |
|---|---|
| P value | 0.8735 |
| Exact or approximate P value? | Exact |
| P value summary | ns |
| Significantly different (P < 0.05)? | No |
| One- or two-tailed P value? | Two-tailed |
| Sum of ranks in column A,B | 2045 , 1960 |
| Mann-Whitney U | 970 |

**Supplementary Figure 8a**

| | KO | KO+WT | KO+C2AmB | KO+C2ABm | KO+C2AmBm |
|---|---|---|---|---|---|
| Number of values | 17 | 24 | 19 | 20 | 19 |
| Mean (pA) | 26.4 | 28.4 | 25.3 | 24.7 | 24.4 |
| Median | 23.5 | 29.5 | 23.8 | 24.8 | 23.5 |
| Std. Deviation | 8.88 | 7.12 | 8.67 | 7.11 | 8.82 |
| Std. Error of Mean | 2.15 | 1.45 | 1.99 | 1.59 | 2.02 |

| Kruskal-Wallis test | |
|---|---|
| P value | 0.4169 |
| Exact or approximate P value? | Approximate |
| P value summary | ns |
| Do the medians vary signif. (P < 0.05)? | No |
| Number of groups | 5 |
| Kruskal-Wallis statistic | 3.920 |
